# *Vibrio campbellii* encodes a distinct set of type III secretion system effectors that mediate cytotoxicity in eukaryotic host models

**DOI:** 10.64898/2026.05.26.727984

**Authors:** Payel Paul, Shir Mass, Hadar Cohen, Matthew L. Bochman, Ram Podicheti, Douglas B. Rusch, Motti Gerlic, Dor Salomon, Julia C. van Kessel

## Abstract

Type III secretion systems (T3SS) are common virulence factors that facilitate the injection of anti-eukaryotic effector toxins that damage and kill target host cells. The sequence and function of the structural and regulatory proteins of these systems are conserved across Gram-negative pathogens, including *Pseudomonas*, *Yersinia,* and *Salmonella.* However, the identity and function of effector proteins are not conserved and are unknown in several relevant human and animal pathogens. Here, we used comparative genomics to identify and characterize the effectors encoded by *Vibrio campbellii* BB120, a crustacean and fish pathogen. We showed that most sequenced *Vibrio* strains belonging to the Harveyi clade, including *V. campbellii*, encode a full set of structural and regulatory genes corresponding to the T3SS1 of *V. parahaemolyticus*; the exception is *V. natriegens*. Transcriptomic and proteomic analyses identified four *V. campbellii* effectors that are secreted by the T3SS. Among these, two effectors are encoded outside the T3SS island, and all effector genes were co-regulated by both the master T3SS regulator ExsA and by the master quorum sensing regulator LuxR. Three effectors – VopS, CopA, and VIBHAR_06684 – exhibited toxic activity in yeast cells or bone marrow-derived macrophages, and the toxicity phenotypes were dependent on a functional T3SS. VopS and CopA are conserved among the queried species of the Harveyi clade. VIBHAR_06684 or VIBHAR_05674 did not show conservation among the queried species. These findings demonstrate that T3SSs in bacteria from the same clade have conserved structural secretion apparatuses but exhibit variance in effector repertoires. We postulate that the functions of effectors differ between species to impart roles in host specificity.

**Author Summary:** Pathogenic *Vibrio* bacteria employ many virulence factors to cause disease in marine animals, including biofilms, proteases, and the secretion of toxins. Here, we identify four secreted toxins encoded by *V. campbellii* that are toxic to eukaryotic cells. These toxins are secreted by the type III secretion system, a needle-like apparatus used by many bacteria to inject toxins into host cells. The toxins identified in this study are regulated by quorum sensing, a method bacteria use to communicate and control virulence genes required for pathogenesis. The results of this study indicate that different *Vibrio* species employ distinct sets of toxins that possibly facilitate their interaction with a host.

## Introduction

*Vibrio campbellii* is a Gram-negative halophilic γ-proteobacterium that belongs to the class *Vibrionaceae* and naturally inhabits the marine environment, specifically the warm waters of Asia, Southern Europe, and South America (1–4). *V. campbellii* belongs to the Harveyi clade, which refers to a cluster of *Vibrio* species that are pathogenic to marine organisms and cause significant economic losses to the fisheries and aquaculture industries around the world (1, 5–7). Gram-negative pathogens such as *Vibrio* species utilize various mechanisms of pathogenesis to invade hosts and establish infection. One of the major mechanisms of pathogenesis employed by pathogens such as *Pseudomonas aeruginosa*, *Yersinia* spp., *Shigella* spp., *Salmonella* spp., *V. parahaemolyticus*, *etc*., is the type III secretion system (T3SS) (8–15). The T3SS is a syringe-like membrane-embedded apparatus, also called the ‘injectisome’, that injects toxic proteins called effectors directly into host cells (8, 9). The effectors target various cellular mechanisms to compromise the host cell machinery and inhibit immune responses (8, 9).

Among *Vibrionaceae* species, the T3SSs in *V. parahaemolyticus* are the most well-studied. *V. parahaemolyticus* encodes two different T3SSs: T3SS1 and T3SS2 (16). While T3SS1 is found in all sequenced clinical and environmental strains of *V. parahaemolyticus*, T3SS2 is only found in clinical isolates (16, 17). T3SS2 mediates enterotoxicity and invasion of host cells and causes gastroenteritis in humans and animal models (18–20). T3SS1 causes lysis of host cells and release of nutrients into the environment, required for the survival of the bacterium (21). *V. campbellii* strains only encode T3SS1, which is the focus of this study; we will refer to it as T3SS throughout this manuscript. Comparative genomics analyses have revealed the presence of T3SS pathogenicity islands in *V. campbellii* strains, including LMB29, 20130629003S01, CAIM 519T, BoB-90, Bob-53, and LA16-V1, which have been isolated either from infected marine organisms or biofouled moorings (2, 3, 22, 23). In addition to these wild isolates, a T3SS is encoded by *V. campbellii* BB120 (*a.k.a.*, ATCC BAA-1116, previously classified as *V. harveyi*) (24), a laboratory strain model organism that also infects brine shrimp in lab infection models (25).

In the well-studied T3SS1 of *V. parahaemolyticus,* there are four known effectors: VopS, VopQ, VopR, and VPA0450 (13, 21). The function and mechanism of action of these effectors are well-characterized. VopS inactivates Rho GTPase family proteins, including Rho, Rac, and Cdc42, by covalently modifying them with an adenosine-5’-monophosphate (AMP) moiety (26). VopS activity leads to the induction of autophagy, cell rounding, and cell lysis (26). VopQ inhibits the fusion of V-ATPase-containing membranes by binding to a conserved V_o_ domain of the V-type H+ ATPase (27). VopR binds phosphatidylinositol-4,5-bisphosphate and localizes to the plasma membrane, causing cell rounding and ultimately cell lysis (28). VPA0450 is an inositol polyphosphate 5-phosphatase that hydrolyzes phosphatidylinositiol-4,5-bisphosphate, leading to blebbing of the plasma membrane, loss of plasma membrane integrity, and ultimately cell lysis (29). Thus, T3SS effectors utilize diverse mechanisms to subvert the host cell machinery and lyse the host cell.

It has been previously shown that *V. campbellii* BB120 encodes a functional T3SS that is active during brine shrimp infection and is regulated by quorum sensing (25, 30–32). *Vibrio* species such as *V. campbellii* and *V. parahaemolyticus* utilize a quorum sensing system (QS) as a mechanism to communicate with bacterial cells within their species, as well as with different species (30, 31, 33). Chemical messengers called autoinducers are secreted by bacterial cells into the external milieu to inform each other about population density (30, 31). Dependent on the population density, either the low cell density regulator AphA is activated, or at high cell density, the regulator LuxR is activated (33). At high cell density, LuxR is maximally expressed and has been shown to repress the T3SS in *V. campbellii* via two mechanisms: 1) transcriptional repression of the T3SS master activator *exsA*, and 2) post-translational repression of ExsA activity through the transcriptional repression of the T3SS positive regulator *exsC* (30, 32). However, the suite of effectors secreted by the *V. campbellii* T3SS is unknown, and thus the mechanisms by which *V. campbellii* uses T3SS to target host cells remain unclear.

In this study, we utilize a multifaceted approach combining genomics, transcriptomics, proteomics, and infection assays to define: 1) the *V. campbellii* T3SS effector repertoire and its regulation by QS, 2) the conditions required by the *V. campbellii* T3SS to infect eukaryotic host models, and 3) the role played by T3SS effectors during host model infection. We found that the *V. campbellii* expresses and secretes at least four T3SS effectors: VopS, CopA, VIBHAR_05674, and VIBHAR_06684. While *copA* and *vopS* are located in the canonical T3SS pathogenicity island on chromosome I, *VIBHAR_06684* and *VIBHAR_05674* are encoded at different loci on chromosome II. Expression in a model eukaryotic cell, the yeast *Saccharomyces cerevisiae*, and infection assays with a eukaryotic immune cell model, murine bone marrow-derived macrophages, revealed the individual and combined toxic roles of the T3SS effectors.

## Results

### T3SS effectors are divergent in Vibrionaceae

Previous studies have shown that the *V. campbellii* T3SS gene operons are highly similar in sequence, organization, and transcriptional regulation to T3SS1 in *V. parahaemolyticus* (31, 32); however, the effectors have not been determined. First, we aimed to determine the extent of T3SS1-like gene conservation among *Vibrio* species using comparative genomics. We analyzed all complete genome sequences for the genus *Vibrio* available through Genbank using *V. parahaemolyticus* RIMD2210633 as the reference strain (Fig. 1, S1). We found that most species belonging to the Harveyi clade, *V. harveyi*, *V. owensiii*, and *V. alginolyticus*, also have a well-conserved T3SS like *V. parahaemolyticus* and *V. campbellii*. Of note, multiple *V. natriegens* strains and a few other species in the Harveyi clade do not appear to encode a T3SS1-like apparatus, which correlates with the finding that *V. natriegens* has not yet been associated with disease in marine organisms. *V. cholerae*, *V. vulnificus*, and *V. coralliilyticus* also do not appear to encode a T3SS1-like apparatus. In *V. parahaemolyticus*, *V. campbellii*, *V. alginolyticus*, *V. owensii*, and *V. harveyi* strains, the proteins predicted to comprise the T3SS structural apparatus are conserved. Conversely, when comparing *Vibrio* strains to *V. parahaemolyticus*, the T3SS effector proteins are highly divergent; *V. campbellii* encodes homologs of the *V. parahaemolyticus* effectors VopQ and VopS, whereas *V. campbellii* does not have homologs of VopR and VPA0450.

**Figure 1.**
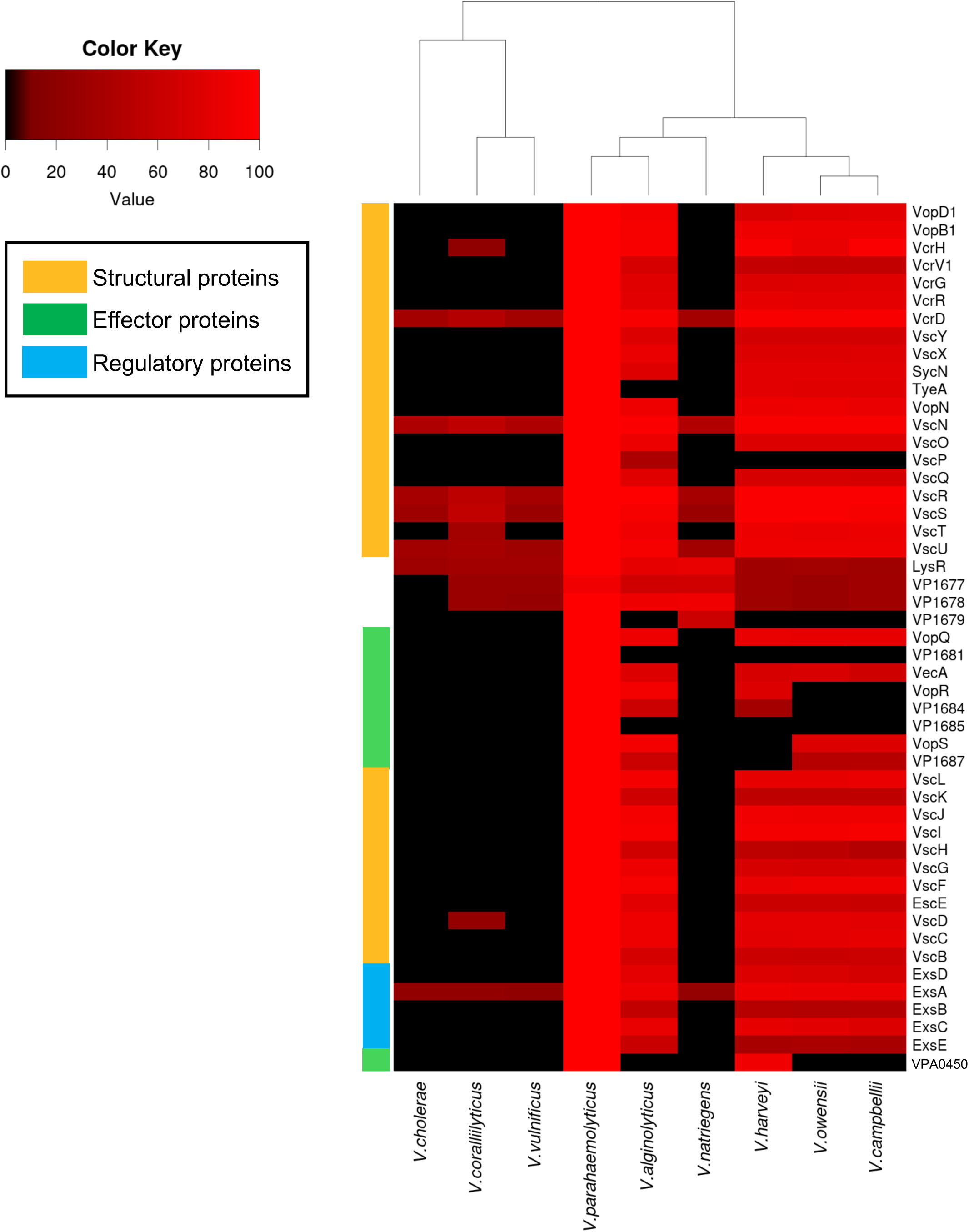
Comparative genomics analysis shows conservation of the T3SS genes in the family *Vibrionaceae*. The percent amino acid identity for each T3SS gene represents the highest scoring homolog across all strains for that species compared to the reference strain *V. parahaemolyticus* RIMD2210633. The dendrogram represents the phylogenetic relationship among the depicted *Vibrio* species.

We observed that the conservation of T3SS structural proteins extends from *Vibrio* species to organisms of distant genera, including *Pseudomonas*, *Yersinia,* and *Salmonella* (Fig. S2), where the vast majority of the genetic, biochemical, and structural biology has been performed to define the T3SS. These conserved structural proteins include the cytoplasmic complex (VscN, VscO, VscL, VscQ, VscK), export apparatus (VscR, VscS, VscT, VscU, LcrD), basal body (VscC, VscD, VscJ), needle complex (VscF, VscI), and the translocon and pore complex (VscE, VscB, VopB1, VopD1, VcrV1) (34–36). We note that although *V. cholerae*, *V. coralliilyticus*, *V. vulnificus*, and *V. natriegens* (among others, Fig. S1) do not appear to encode a functional T3SS, they do encode homologs of certain structural proteins, including VscN, LcrD, VscR, VscS, VscT, and VscU. This could indicate conservation of structure between the T3SS apparatus and the flagellum, with the two nanomachines being known to share a common ancestor (37, 38). VscR, VscS, VscT, LcrD, and VscU are part of the export apparatus and are homologous to flagellar export apparatus proteins FliP, FliQ, FliR, FlhA, and FlhB, respectively (38). VscN is an ATPase and is homologous to the flagellar ATPase protein FliL. Conversely, ExsA and LysR are transcription regulators with widely distributed homologs across the genus. From these comparative genomics studies, we conclude that species of the Harveyi clade (*V. parahaemolyticus*, *V. campbellii*, *V. harveyi*, *V. alginolyticus*, *V. owensii*) encode T3SSs that have the potential to form fully functional T3SS machines, whereas *V. cholerae*, *V. coralliilyticus*, *V. vulnificus*, and *V. natriegens* likely do not encode a T3SS1-like apparatus. We also conclude that the structural T3SS genes identified and studied in other γ-proteobacteria are conserved in *V. campbellii*, whereas the putative effector proteins encoded within and outside the T3SS gene cluster differ between *Vibrio* species.

### ExsA and LuxR co-regulate the expression of orphan T3SS effectors

Previous studies reported *V. campbellii* BB120 LuxR as an inhibitor of T3SS gene expression at high cell density through the repression of the T3SS activator ExsA (30, 32, 33). As in other T3SSs, including *Pseudomonas, Yersinia,* and *V. parahaemolyticus*, the conserved ExsA transcription factor is an AraC-type regulator that activates transcription of the suite of genes encoding the T3SS structural component and effectors (39–42). In *V. campbellii*, LuxR is a broad co-regulator of the T3SS genes, known to repress 49 genes in the T3SS pathogenicity island on chromosome I of *V. campbellii* BB120 (30–33). We hypothesized that T3SS effectors encoded outside of the pathogenicity island are also regulated by ExsA and LuxR. To identify the complete co-regulon of these two master transcription factors, we performed RNA-seq comparing transcripts of cells collected at high cell density (OD_600_ = ∼1.0) from *V. campbellii* wild-type, Δ*luxR*, Δ*exsA,* and Δ*luxR* Δ*exsA* strains (Dataset S1). The results showed that ExsA is indeed necessary for the transcriptional activation of 40 of the 49 genes previously reported to comprise the T3SS pathogenicity island on chromosome I (Fig. 2A, Dataset S1); these genes are also repressed by LuxR. However, 9 of the 49 genes previously thought to be regulated by ExsA and the QS system did not adhere to the predicted regulatory pattern (*VIBHAR_01714-01722*). Further inspection of these genes revealed that they contain transposase sequences (Fig. 2A, Dataset S1).

**Figure 2.**
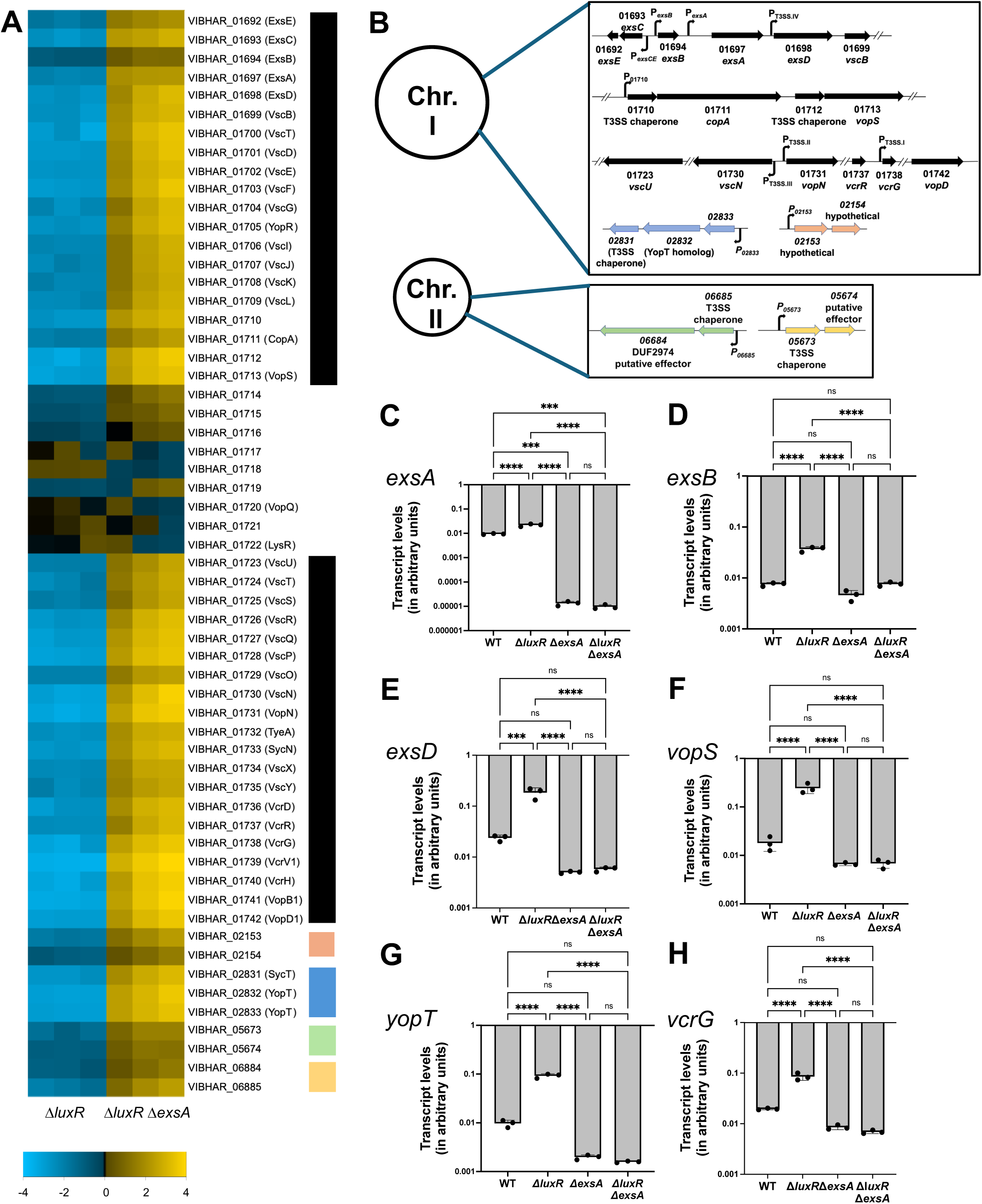
RNA sequencing analysis of genes activated by ExsA and repressed by LuxR at high cell density in *V. campbellii* BB120. **A.** Heat map shows expression levels of T3SS genes across strains. Colored bars correspond to operons in chromosomes I and II depicted in the next figure. **B.** Organization of T3SS genes into four structural operons in both chromosomes. Genes depicted in black are part of the canonical T3SS pathogenicity island. Genes depicted in multi-color represent new-found operons regulated by ExsA and LuxR and predicted to be part of the T3SS. **C-H.** RT-qPCR data with isogenic BB120 strains to measure absolute transcript levels of *exsA, exsB, exsD, vopS, yopT, and vcrG* genes compared to internal control *hfq*. Error bars represent the standard deviation of the mean for triplicate biological replicates. A one-way analysis of variance (ANOVA) test was performed on normally distributed data (Shapiro-Wilk test) followed by Tukey’s multiple comparisons test. (*p*<0.0001, *n*=3)

Our RNA-seq assay also revealed that ExsA and LuxR co-regulate additional operons in separate locations on chromosomes I and II. These include *VIBHAR_02831-02833* and *VIBHAR_02153-02154* in chromosome I, and *VIBHAR_05673-05674* and *VIBHAR_06684-06685* in chromosome II (Fig. 2A-B). Among the genes found to be co-regulated by ExsA and LuxR are those predicted to encode effector homologs of *V. parahaemolyticus* VopS and *Y. pestis* YopT. We also identified genes encoding putative T3SS effectors *VIBHAR*_*01711*, *VIBHAR_05674,* and *VIBHAR_06684*, based on their proximity to predicted chaperone-encoding genes. The RNA-seq data were validated through RT-qPCR, corroborating the observed transcriptional regulation by LuxR and ExsA at high cell density (Fig. 2C-H). From these results, we conclude that ExsA and LuxR co-regulate 49 genes, and we hypothesize that these are associated with T3SS regulation and function. We also conclude that there are potential effectors regulated by ExsA and LuxR outside the main pathogenicity island on chromosome I.

### Mg^2+^ induces T3SS activity in *V. campbellii*

A previous examination of the *V. campbellii* BB120 T3SS showed that this system is functional and secretes at least one known secreted protein, VopD (31, 33). VopD and VopB are the type III translocon subunits comprising the tip of the type III secretion channel, forming a pore at the host cell membrane for targeted injection of the effectors (38). Thus, we monitored VopD secretion as an indicator of T3SS activity, to determine the optimal conditions for type III-mediated secretion in *V. campbellii*. As a negative control in which the T3SS is inactive, we constructed a strain lacking *vscC*, a gene encoding a key T3SS structural protein (43, 44). VscC is a homolog of the SctC protein, which forms the upper ring of the basal body in the T3SS apparatus (35, 38). In *V. parahaemolyticus*, the *vscC* gene is essential for enterotoxicity against the host (45). We constructed the *vscC* deletion in the Δ*luxR* background, reasoning that the T3SS genes would be transcriptionally activated, but no secretion would occur. We collected whole cell lysates and cellular supernatants from the Δ*luxR* and Δ*luxR* Δ*vscC* strains and performed immunoblot assays to monitor VopD expression and secretion. To this end, we examined several conditions because previous studies in *V. parahaemolyticus* and *P. aeruginosa* showed that the presence of calcium in growth media acts as a repressor of type III secretion, whereas chelation of calcium from growth media using EGTA induces type III secretion (45–47). We tested the following conditions using our standard LB Marine media (LM; Lysogeny Broth with 2% [w/v] NaCl): 1) LM only, 2) LM + 15 mM MgSO_4_, 3) LM + 5 mM EGTA, 4) LM + 15 mM MgSO_4_ + 5 mM EGTA, and 5) LM + 15 mM CaCl_2_. VopD was secreted via T3SS under all media conditions tested except when EGTA was added to the growth media (Fig. 3, S3), indicating that cation chelation blocks T3SS-mediated secretion of VopD. The addition of excess calcium did not affect VopD secretion. However, the addition of magnesium (MgSO_4_) elevated both VopD expression and secretion (Fig. 3, S3). In a previous study, we had also observed that the addition of 15 mM MgSO_4_ to LM led to higher transcription of T3SS genes than growth in LM alone (32). The results indicate that deletion of QS regulation (Δ*luxR*) is sufficient to activate T3SS expression and secretion. Deletion of *vscC* abolished VopD secretion but did not dramatically alter expression levels. Additionally, magnesium may play an important role in the expression and secretion of T3SS genes. Our attempt at complementing *vscC* ectopically was unsuccessful in restoring secretion (Fig. S4). The likely reason is that deleting *vscC* has a polar effect on the expression of the flanking structural genes *vscB* (VIBHAR_01699) and *vscD* (VIBHAR_01701).

**Figure 3.**
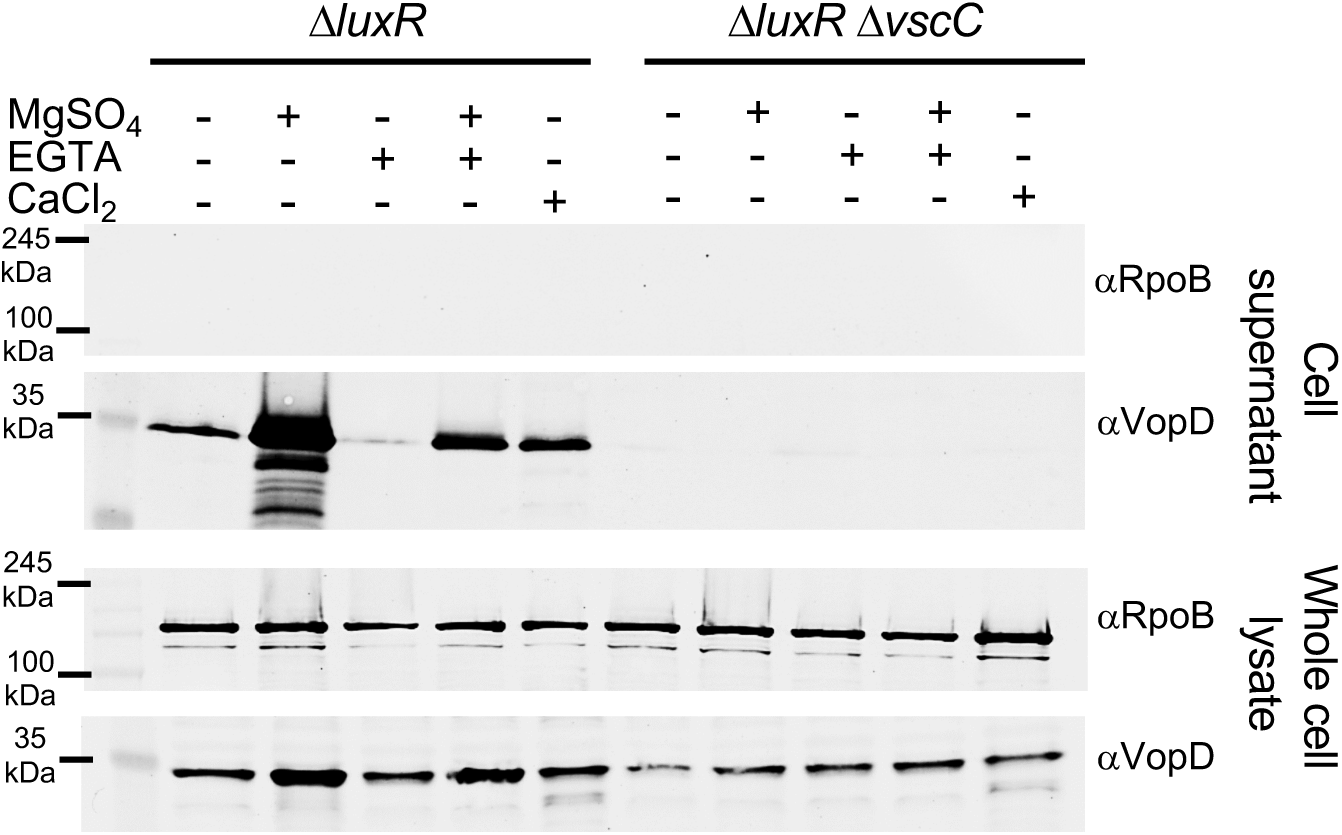
Mg induces secretion of VopD. Secretion assays followed by western blot on cellular supernatants and whole cell lysates in Δ*luxR* and Δ*luxR ΔvscC* strains under different media conditions including LM only, LM+15 mM MgSO_4_, LM+5 mM EGTA, LM+15 mM MgSO_4_+5 mM EGTA, and LM+15 mM CaCl_2_. RNA polymerase β subunit (Rpoβ) was used as a loading control. VopD was blotted with anti-VopD antibodies. Rpoβ was blotted with anti-Rpoβ antibodies.

### *V. campbellii* secretes four putative T3SS effectors

After identifying an optimal media condition that induces type III-mediated secretion in *V. campbellii*, we next sought to determine what proteins are secreted specifically by the T3SS and define its ‘secretome’. To this end, we collected cellular supernatant from the T3SS^+^ strain (Δ*luxR*) and T3SS^−^ strain (Δ*luxR* Δ*vscC*) and performed mass spectrometry to compare protein secretion (Dataset S2). The results revealed the presence of a suite of secreted proteins specific to the T3SS^+^ strain; the 10 proteins that were enriched (> 2-fold label-free quantification intensity and *p* < 0.05) in the Δ*luxR* compared to Δ*luxR* Δ*vscC* supernatant are encoded within the T3SS pathogenicity island or in one of the orphan operons identified by RNA-seq as co-regulated by ExsA and LuxR (Fig. 4A, 2A-B, Dataset S2). Among these are T3SS regulatory proteins ExsE and VopN, type III needle protein VscF, V antigen VcrV, and the type III translocons VopB and VopD (Fig. 4A, Dataset S2) (35, 37, 38, 48). We predict that the remaining four proteins, for which we confirmed T3SS-mediated secretion using immunoblot assays when expressed from a plasmid (Fig. 4B, S5), are T3SS effectors:

**Figure 4.**
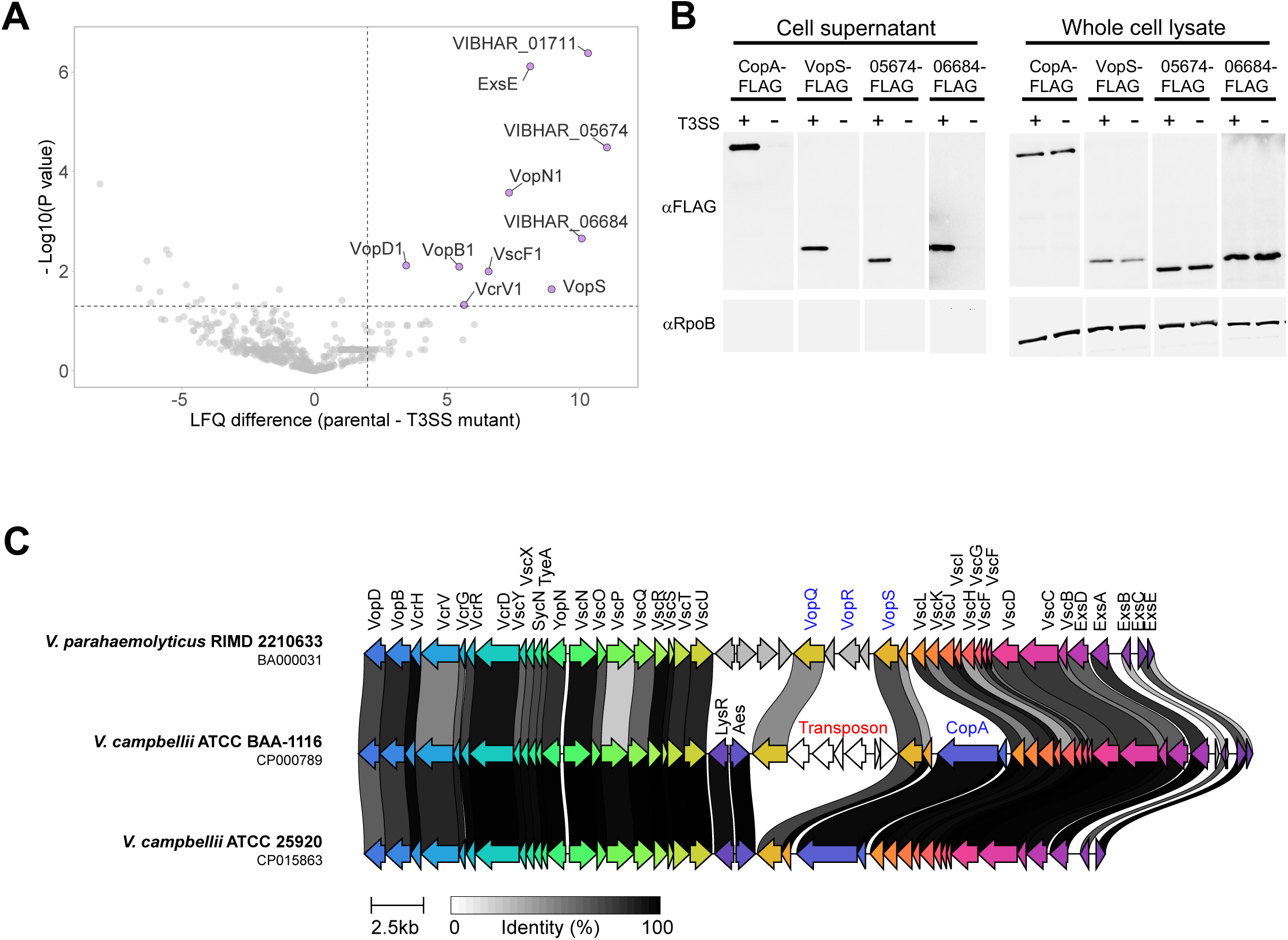
*V. campbellii* BB120 T3SS secretome. **A.** A volcano plot summarizing the comparative analysis of proteins identified in the media of *V. campbellii* BB120 T3SS^+^ (Δ*luxR*) and T3SS^−^ (Δ*luxR* Δ*vscC*) strains, using label-free quantification (LFQ). The average LFQ signal intensity difference between the T3SS^+^ and T3SS^−^ strains is plotted against the −Log10 of Student t-test P-values (*n* = 3 biological replicates). Proteins that were significantly more abundant in the secretome of the T3SS^+^ strain (difference in average LFQ intensities > 2; *P* value < 0.05) are denoted in purple. **B.** Secretion assay followed by western blot with T3SS^+^ (Δ*luxR*) and T3SS^−^ (Δ*luxR* Δ*vscC*) strains with the whole cell lysate and cell supernatant. The putative effector proteins were FLAG-tagged in the C-terminus and overexpressed from an ectopic plasmid in each strain. P*_tactheo_*-*copA-FLAG* (pPP67) was induced with 100 μM IPTG and 10 μM theophylline, P*_tactheo_*-*vopS-FLAG* (pPP68), P*_tactheo_-VIBHAR_05674-FLAG* (pPP71) were induced with 10 μM IPTG and 1000 μM theophylline, and *P_tactheo_-VIBHAR_06684-FLAG* (pPP72) was induced with 100 μM IPTG and 1000 μM theophylline. FLAG-tagged proteins were blotted with anti-FLAG antibody. RNA polymerase β subunit (Rpoβ) was used as loading control and blotted with anti-Rpoβ antibody. Strains expressing *copA, vopS,* and *VIBHAR_05674* were grown in LM medium with 15 mM MgSO_4_. The strain expressing *VIBHAR_06684* was grown in LM medium with 15mM MgSO_4_ and 5 mM EGTA. **C.** Genome names and accession numbers are denoted. Coding sequences are represented by arrows and colored to reflect homologous groups identified by Clinker. Connecting shaded grey rectangles represent the percentage of amino acid identity. Predicted effectors are denoted in blue.

#### VIBHAR_01713 (WP_012127534.1)

Found within the T3SS cluster, this gene encodes a homolog of *V. parahaemolyticus* VopS (26), which covalently modifies Rho family GTPases, including Rac, Rho, and Cdc42, through ‘AMPylation’, which is the transfer of adenosine 5’ monophosphate onto a target residue. We will refer to VIBHAR_01713 as VopS for the remainder of this manuscript. The *V. campbellii* BB120 VopS shares 74% amino acid sequence homology with VopS from *V. parahaemolyticus*.

#### VIBHAR_01711 (WP_012127533.1)

Found within the T3SS cluster, this gene encodes a potential effector with a conserved ADP-ribosyl transferase domain. *P. aeruginosa* also has T3SS effectors with actin-ADP ribosylation activity called ExoS and ExoT (49–51). They are known to modify Rho-family GTPases through ADP-ribosylation. However, the target for CopA in *V. campbellii* BB120 is unknown. We name this *V. campbellii* effector CopA (*V. <u>c</u>ampbellii* <u>o</u>uter <u>p</u>rotein <u>A</u>DP-ribosyltransferase).

#### VIBHAR_06684 (WP_012130081.1)

Found outside the T3SS cluster, this gene is adjacent to a predicted T3SS chaperone. The Protein is predicted to contain a C-terminal alpha/beta hydrolase family domain, according to the NCBI Conserved Domain Database (52).

#### VIBHAR_05674 (WP_041853546.1)

Found outside the T3SS cluster, this gene is adjacent to a predicted T3SS chaperone. Although this protein was identified as secreted, no function has yet been predicted for VIBHAR_05674 based on sequence and conserved domain analyses.

We next evaluated why the secretome analysis failed to identify putative homologs of known effectors that we annotated in the *V. campbellii* BB120 genome (*i.e*., VopQ and YopT). Analysis of the BB120 VopQ homolog, encoded within the T3SS cluster, revealed that the *vopQ* gene is missing the N-terminal sequence corresponding to amino acids 30-101 in the *V. parahaemolyticus* homolog, and that it is fused with an upstream chaperone gene (Fig. 4C). Thus, we hypothesize that the BB120 VopQ homolog is non-functional. The *Y. pestis* YopT homolog encoded by *V. campbellii* did not appear in the secretome data either. Analysis of the *yopT* gene sequence revealed a premature stop codon, splitting the gene into two loci, thus likely rendering it non-functional.

### Three *V. campbellii* effectors are toxic in yeast

To determine whether the four putative effectors have toxic effects in eukaryotic cells, we used yeast as a model eukaryotic cell to monitor the effect of effector expression on cell viability (28). To this end, we constructed yeast strains expressing individual genes from the *GAL1* galactose-inducible promoter and grew the cells either in the presence of 2% (w/v) glucose (repressing conditions) or 2% (w/v) galactose (inducing conditions) (Fig. 5A). We observed that VIBHAR_05674 did not show any toxicity in yeast (Fig. 5A). Conversely, galactose induction of VIBHAR_06684 led to reduced fitness of yeast cells compared to the empty vector control (Fig. 5A). We note that we were unsuccessful in transforming the plasmids expressing CopA and VopS into yeast cells; we show that no transformants were obtained in triplicate transformation assays compared with expression plasmids for VIBHAR_05674 and VIBHAR_06684 (Fig. 5B). We hypothesized that CopA and VopS were exceedingly toxic to yeast cells even under non-inducing conditions due to leaky transcription from the *GAL1* promoter (53), leading to their death during transformation. To test our hypothesis, we identified putative active sites based on previous studies in *V. parahaemolyticus* VopS and *P. aeruginosa* ExoS (26, 49). We constructed mutant alleles at the predicted active sites of the two genes: *vopS* H347A and *copA* E836D/E838D. Yeast transformants were obtained with the active site mutants of CopA and VopS at similar levels to the empty vector (Fig. 5B). We interpret these results to mean that active CopA and VopS proteins are lethal to yeast cells. From these data, we conclude that *V. campbellii* BB120 encodes three proteins that target a conserved eukaryotic target found in yeast: CopA, VopS, and VIBHAR_06684.

**Figure 5.**
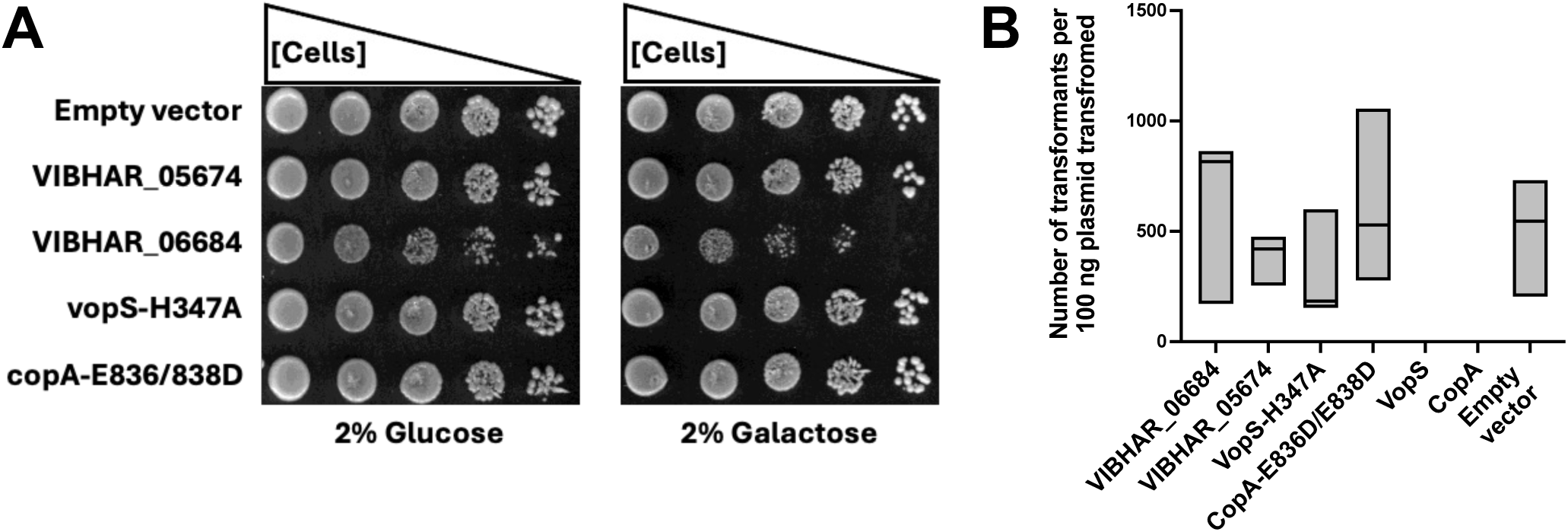
Toxicity assays conducted in *Saccharomyces cerevisiae* with *V. campbellii* BB120 effectors transformed into vectors. **A.** Wild-type *Saccharomyces cerevisiae* cells were transformed with empty vector or plasmids encoding the indicated genes under the control of the glucose-inhibited, galactose-inducible *GAL1* promoter. Transformed yeast cells were plated on the inhibitory 2% glucose media plates or the inducing 2% galactose media plates. **B.** The number of transformed colonies for each vector transformed carrying a *V. campbellii* BB120 effector gene was plotted into a bar graph to display transformation efficiency. The graphed bars span the minimum and maximum of the colony counts from three independent experiments, with the horizontal lines representing the median values. No values are indicated for VopS or CopA because no colonies were recovered.

### *V. campbellii* T3SS induces macrophage cell death

To determine whether the T3SS is employed to intoxicate host cells, we monitored infection of murine bone marrow–derived macrophages (BMDMs), which served as a model host immune cell. The addition of the parental *V. campbellii* T3SS^+^ strain (Δ*luxR*) to BMDMs induced cytotoxicity in ∼15% of the cells within 160 minutes (Fig. 6A, 6B). This cytotoxicity is T3SS-dependent, as no cell death was observed when an isogenic T3SS^−^ strain (Δ*luxR* Δ*vscC*) was used.

**Figure 6.**
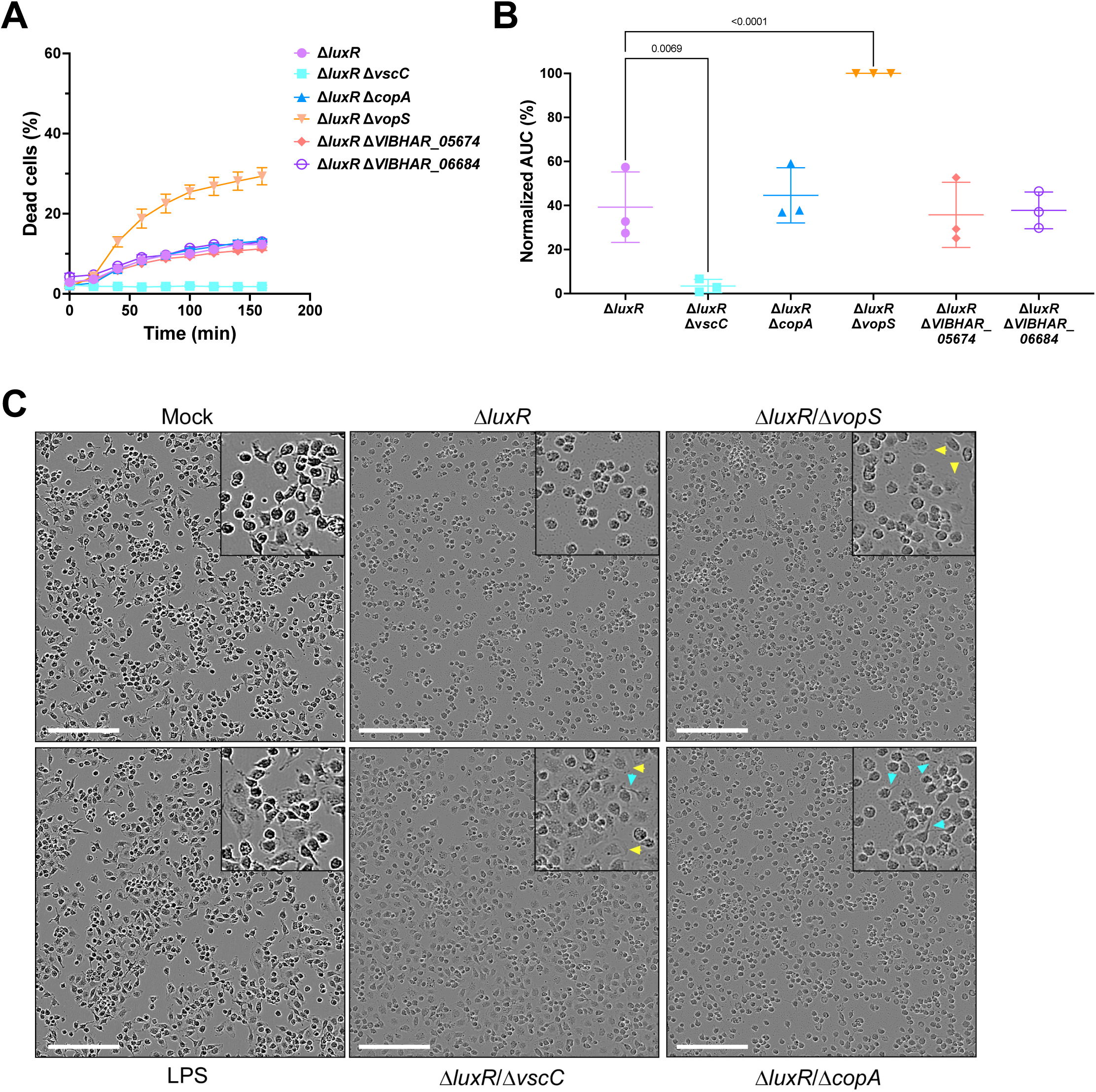
The *V. campbellii* T3SS induces cytotoxicity in host macrophages. **A-B.** Assessment of cell death upon infection of bone marrow-derived macrophages (BMDMs) with the indicated *V. campbellii* strains. Approximately 3.5×10^4^ BMDMs were seeded into 96-well plates in triplicate and were infected with *V. campbellii* strains at a multiplicity of infection (MOI) of 5. Propidium iodide (PI) was added to the medium prior to infection, and its uptake kinetics were assessed during 160 minutes using real-time microscopy (Incucyte SX5) (A); a representative result is shown; n = 3 technical repeats). BMDM cell death was analyzed as the area under the curve (AUC) across 3 independent experiments and normalized to the relative AUC from infection with the *vopS* mutant, set at 100% (B). The data are presented as the mean ± SD. Statistical comparisons in (B) were performed using a one-way ANOVA, followed by Tukey’s multiple comparison test. Significant *P* values (<0.05), compared to the parental T3SS^+^ strain, are denoted. **C.** Representative microscopy images of BMDMs, 1h post-infection with the indicated *V. campbellii* strains. The squares on the top right corner of each image are a 2x magnification of a random area from the image. Yellow arrows denote examples of flat cells; cyan arrows denote examples of cell extensions. Mock = uninfected. Scale bar = 200 µm.

Next, we sought to investigate the possible contribution of individual T3SS effectors to cytotoxicity. Although the deletion of either *copA*, *VIBHAR_05674*, or *VIBHAR_06684* did not affect the T3SS-dependent cytotoxicity of *V. campbellii*, the deletion of *vopS* led to enhanced cytotoxicity, reaching ∼30% within 160 minutes of infection (Fig. 6A, 6B). These findings are consistent with a previous report showing that the VopS and VopQ homologs from *V. parahaemolyticus* suppress inflammasome activation by interfering with NLRC4-induced cell death, thereby evading host pro-inflammatory responses (54). Thus, deletion of *vopS* and *vopQ* increased T3SS-dependent caspase activation and cell death (pyroptosis). Notably, as mentioned above, *V. campbellii* BB120 lacks a functional secreted VopQ homolog, thus the deletion of *vopS* is likely sufficient to eliminate the suppression of pyroptosis.

In addition, we observed distinct morphological changes in BMDMs during infection with the effector-deletion mutant strains. Infection with the parental T3SS^+^ strain induced pronounced cell rounding within one hour, whereas most of the cells infected with the T3SS-inactive mutant were flat and exhibited prominent extensions. Infection with the *copA* mutant maintained cells with cellular extensions, whereas infection with the *vopS* mutant maintained a population of flat cells (Fig. 6C). Notably, the effect of *vopS* deletion on BMDM morphology was to be expected considering that VopS homologs target Rho GTPases and affect the actin cytoskeleton (26).

VopD secretion assays were performed on the *vopS* and *copA* effector deletion strains to confirm that the phenotypes observed during BMDM infections were not due to changes in T3SS secretion potential (Fig. S6). Indeed, we observed no change in VopD secretion, suggesting that the observed phenotypes result from changes in specific effector activities.

### The effector suite of *V. campbellii* BB120 is unique

We examined the conservation of the four *V. campbellii* BB120 effectors across the *Vibrio* phylogenetic tree (Fig. 7). We observed that VIBHAR_05674 and VIBHAR_06684 are not conserved even among the other *V. campbellii* strains that we queried (complete genomes in Genbank). However, NCBI protein BLAST searches revealed that VIBHAR_05674 has a narrow phylogenetic distribution with presumed homologs in two species, including *V. caribbeanicus* and *Algicola sagamiensis*. VIBHAR_06684, on the other hand, has a wider phylogenetic distribution with presumed homologs in *V. sagamiensis*, *V. aestuarinus*, *V. misgurnus*, *Aeromonas schubertii*, and *Edwardsiella ictaluri*. Of note, many of these bacterial species were isolated from diseased fish and other marine organisms (55–59). Conversely, VopS and CopA appear to be conserved in the queried *V. campbellii* species and a few other strains, including *V. owensii*, *V. harveyi*, and *V. jasicida*. Most strains in the Harveyi clade encode a VopS homolog (Fig. 7). Among the strains that were predicted to encode a T3SS structure (100), only 3 did not encode a VopS homolog. These results indicate that the *V. campbellii* effector repertoire has both conserved and diverse effectors among the known types of effectors in the *Vibrio* genus.

**Figure 7.**
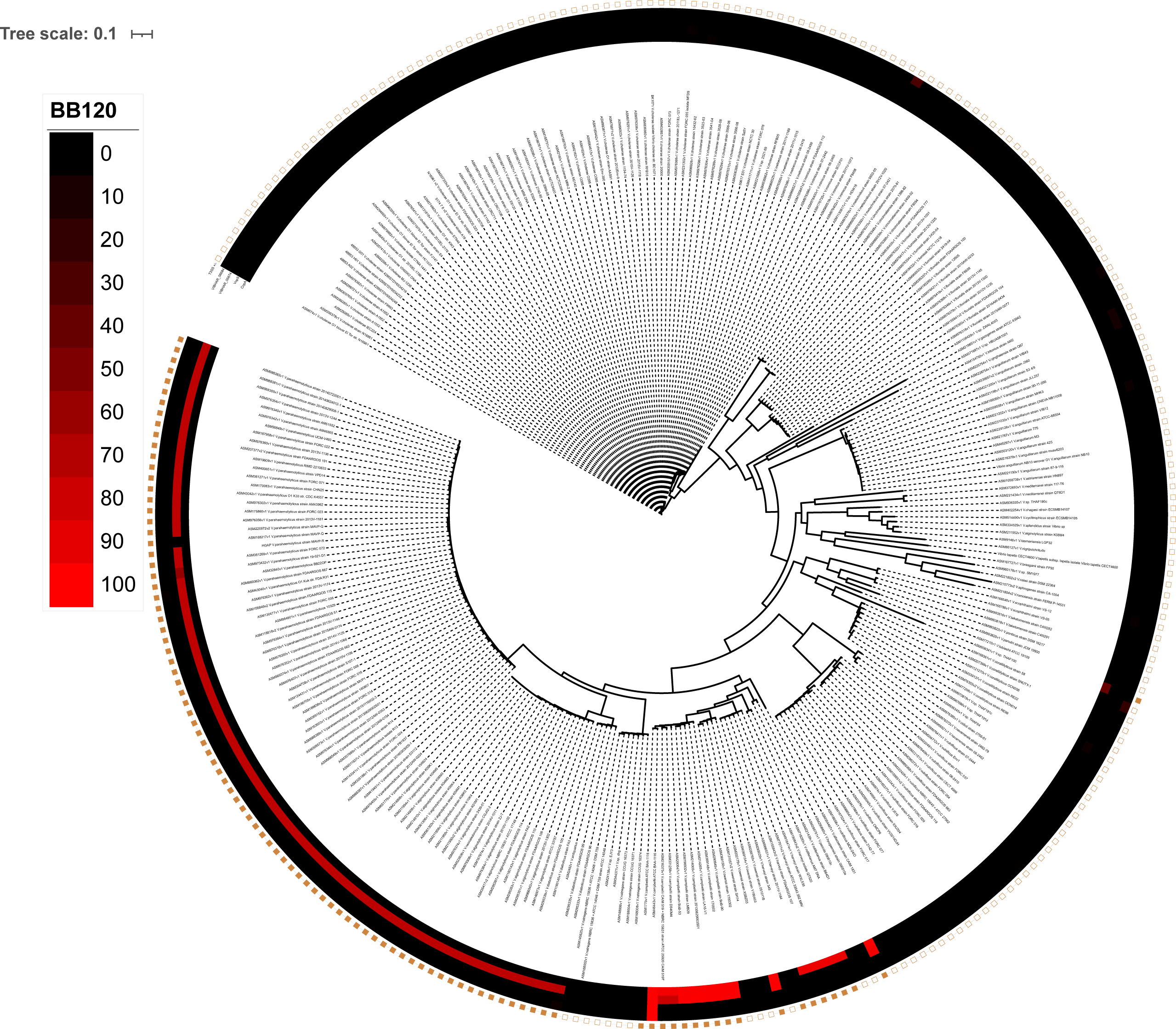
Comparative genomics analysis shows the gene conservation in the family *Vibrionaceae*. *Vibrio campbellii* BB120 has been used as the reference strain for the comparative genomics analysis. The presence of a T3SS (+/−) was determined based on the data presented in Figure S1. Strains that were predicted to have a T3SS are designated as “+” and have a filled square.

## Discussion

In the *Vibrio* genus, each species has distinct host niches in the marine environment, yet it is not well understood what factors determine host compatibility for each bacterium, nor what makes colonization or infection effective in each species. Here, we focused on the T3SS system of *V. campbellii*, which is a widespread burden on the aquaculture industry (60). Using known T3SS genes from *V. parahaemolyticus* as queries, we determined the conservation of T3SS components in *V. campbellii* and other *Vibrio* species. Gene clusters containing these genes were found in *Vibrio* species belonging to the Harveyi clade, including *V. campbellii.* Our transcriptomics and proteomics analyses revealed notable co-regulation of *V. campbellii* T3SS genes, both within the pathogenicity island and outside this region. We determined that *V. campbellii* T3SS secretes at least four effectors, three of which are toxic when ectopically expressed in a model eukaryotic cell: CopA, VopS, and VIBHAR_06684. CopA and VopS (VIBHAR_01711 and VIBHAR_01713, respectively) are encoded by genes present within the T3SS pathogenicity island on chromosome I. However, putative effector VIBHAR_06684 is encoded on chromosome II in a divergent location away from the known T3SS pathogenicity island. This is reminiscent of the finding that VPA0450, a T3SS effector in *V. parahaemolyticus*, is located outside the pathogenicity island and on chromosome II (61). The field’s knowledge of homologs in other organisms allows us to postulate their functions. We hypothesize that CopA is an ADP-ribosylating protein that targets Rho GTPase family proteins, VopS is an AMPylating protein that also targets Rho GTPases, and VIBHAR_06684 likely encodes an alpha/beta hydrolase. Future studies of these three proteins will determine their mode of action, target, and level of activity in host cells.

We used two eukaryotic cell models, yeast (*Saccharomyces cerevisiae*) and murine bone marrow–derived macrophages (BMDMs) to assay for toxicity of the four *V. campbellii* T3SS effectors. Ectopic expression in *S. cerevisiae* showed that VIBHAR_06684 is cytotoxic. Cells into which DNA was introduced to express CopA and VopS did not even survive the transformation. However, transformed cells were obtained with constructs expressing mutant alleles that inactivated the putative catalytic sites of the toxins. This could be indicative of the highly toxic effects of CopA and VopS in eukaryotic cells. However, further testing is required to determine their effect on higher organisms. The function of VIBHAR_05674 will have to be studied further to understand its role in T3SS-mediated pathogenicity. It will be necessary to determine which effectors are necessary and which are sufficient for causing disease in natural hosts.

The many elements that dictate host specificity among *Vibrio* species include broadly important factors such as nutrient acquisition and utilization, survival under stressful conditions (temperature, salinity, pH, *etc.*), use of surface adhesion and motility pili, and acquisition of resistance traits via horizontal gene transfer (62–66). We speculate that secretion of host-specific targeting enzymes via T3SSs also plays an important role in tissue and host specificity. To date, the T3SSs of *V. parahaemolyticus* are the most well-studied of the *Vibrio* species, and the effectors secreted by both its T3SS1 and T3SS2 are known to target and modulate host cell viability. *V. campbellii* BB120 appears to have lost functional alleles of two effectors homologous to effectors found in other pathogens – VopQ and YopT – through deleterious mutations. Thus, *V. campbellii* BB120 only shares a single known effector with *V. parahaemolyticus* – VopS – but has shared effectors with other *V. campbellii* strains and a few other species. The diversity of effector repertoires is hypothesized to reflect two groups: conserved ‘core’ effectors enabling common infection mechanisms (*e.g.*, VopS), and distinct, divergent effectors that may enable more specified host targeting functions (*e.g.*, CopA). The recovery of known and novel effectors from the secretome analysis indicates that this methodology could be extended to other *V. campbellii* strains to identify conserved core effectors and divergent effectors. Further studies on the diversity and functions of T3SS effectors in *Vibrio* species may help distinguish these two groups and assess their contributions to host specificity.

## Materials and Methods

### Bacterial strains and media

All strains used in this study are listed in Tables S1 and S2. The *Escherichia coli* S17-1 λ−*pir* strain was used for cloning purposes. *E. coli* strains were cultured at 37°C with shaking (250–275 RPM) in Lysogeny Broth (LB) media with 100 μg*/*mL kanamycin, 100 μg*/*mL gentamycin, and*/*or 10 μg*/*mL chloramphenicol when selection was required. *V. harveyi* BB120 was recently reclassified as *Vibrio campbellii* BB120 (a.k.a., ATCC BAA-1116) (67). BB120 and derivatives were cultured at 30°C with shaking (250–275 RPM) in LB Marine (LM) medium with 100 μg*/*mL kanamycin, 10 μg*/*mL chloramphenicol, and*/*or 50 μg*/*mL polymyxin B when selection was required. Plasmids were transformed into electrocompetent *E. coli* S17-1 λ*pir* cells and subsequently conjugated into *V. campbellii* strains. *V. campbellii* exconjugants were selected using polymyxin B (50 U*/*mL). In case of strains transformed with inducible plasmid pPP25 and pPP62, 10 μM IPTG, and 10 μM, 100 μM, and 100 μM theophylline were used to induce expression from the plasmids.

### Molecular and chemical methods

PCR was performed using Phusion HF polymerase (New England Biolabs) and Iproof HF polymerase (BioRad). All oligonucleotides were ordered from Integrated DNA Technologies (IDT). PCR products and plasmids were sequenced using Eurofins Genomics. Cloning procedures are available upon request. DNA samples were resolved using 1% (w/v) agarose (1× TBE). Unless otherwise noted, data are plotted for triplicate independent experiments. Symbols on graphs represent the mean values, and error bars are standard deviations. Statistical analyses were performed with GraphPad Prism version 10.2.0. Additional information about statistical analyses is included in the figure legends.

### Construction of deletion/epitope-tagged strains

All *V. campbellii* BB120 derivative strains in this study were constructed following a previously published technique (68). Briefly, the pRE112 suicide vector was used to construct unmarked deletions or insertions of epitopes in which 1000 bp of upstream and downstream flanking sequence was cloned into pRE112. The pRE112 derivatives were conjugated into *V. campbellii* and selected on chloramphenicol to induce chromosomal recombination of the plasmid. Subsequently, the plasmid was excised via counterselection on 15% (w/v) sucrose. Cells in which the plasmid excision yielded a non-WT locus were detected via colony PCR. All gene deletions were confirmed by DNA sequencing through Eurofins.

### RT-qPCR, RNA extraction, RNA-seq data analysis

Strains were inoculated in 5 mL LM and grown overnight shaking at 30°C at 275 RPM. Each strain was back-diluted 1:1000 in LM and 15 mM MgSO_4_ and grown shaking at 30°C at 275 RPM until they reached an OD_600_ =□1.0. 2 mL cells were collected by centrifugation at 3700 RPM at 4°C for 10 min, the supernatant was removed, and the cell pellets were flash frozen in liquid N_2_ and stored at −80°C. RNA was isolated from pellets using a TRIzol/chloroform extraction protocol and treated with DNase via the DNA-free™ DNA Removal Kit (Invitrogen) as previously described (69). Quantitative reverse transcriptase real-time PCR (RT-qPCR) was used to quantify transcript levels of T3SS genes in different regulatory conditions and was performed using the SensiFast SYBR Hi-ROX One-Step Kit (Bioline) according to the manufacturer’s guidelines. Primers were designed to have the following parameters: amplicon size of 100 bp, primer size of 20–28□bases, and melting temperature of 55–60°C. All reactions were performed using a Step One Real PCR system with 0.4 μM of each primer and 200 ng of template RNA (20 μL total volume). All RT-qPCR experiments were normalized to the internal standard *hfq* gene. The ΔΔ*C_T_* or standard curve methods were used to analyze data from at least three independent biological replicates with two technical replicates each.

RNA-seq was performed at the Center for Genomics and Bioinformatics at Indiana University. Sequenced reads were processed for adapter removal and quality filtering using Trimmomatic (v0.38) (70). Filtering required an average base quality score ≥20 within a sliding window of three bases and retained only sequences trimmed to a minimum length of 20 bases (parameters: ILLUMINACLIP:adapters.fa:2:20:7 LEADING:20 TRAILING:20 SLIDINGWINDOW:3:20 MINLEN:20). Filtered read pairs were aligned to the *Vibrio campbellii* ATCC BAA-1116 (BB120) genome sequence (GenBank accessions CP000789.1, CP000790.1, and CP000791.1) using Bowtie2 (v2.4.2) (71). Read pairs that uniquely and concordantly mapped to annotated genes were quantified with feature Counts (v2.0.0) (72) from the Subread package (parameters: -t gene -g locus_tag -s 2 -p -B -C). Differential expression analysis comparing Δ*exsA* and Δ*luxR* mutants against wild type was performed using DESeq2 (v1.24.0) (73). Regularized log-transformed expression data from Δ*luxR* and Δ*luxR* Δ*exsA* replicate libraries were represented in a heatmap using hierarchical clustering.

### Secretion assays

Strains were inoculated in 5 mL LM and grown overnight shaking at 30°C at 275 RPM. Each strain was back-diluted 1:100 in LM+ 15 mM MgSO_4_ + 5 mM EGTA (unless noted otherwise), standardized to OD6_00_ 0.05, and grown with shaking at 30°C at 275 RPM until they reached an OD_600_ ==:□1.0. 1 mL of the culture was pelleted at 13000 rpm for 3 minutes and resuspended in lysis buffer containing 1mL Bugbuster, 5μL 10 mg/mL lysozyme, and 1μL Benzonase. The lysed cells were resuspended in 2X SDS PAGE sample buffer and boiled at 95°C for 5 minutes. 15μL of each sample was loaded onto a 12.5% SDS-PAGE gel and run at 120 V for 70 minutes. The rest of the culture was pelleted down at 4000 rpm at 4°C for 10 minutes. The supernatant was collected by filtering through a 0.22 μM PVDF filter. The supernatant was then treated with deoxycholate to a final concentration of 150 μg/mL. Ice cold trichloroacetic acid (TCA) was then added to a final concentration of 8% (v/v) and the supernatant was incubated overnight at 4°C on ice. On the following day, the supernatant was washed with ice cold acetone twice, followed by neutralization with 100 mM Tris at pH 8.0. The samples were treated with 2X SDS PAGE sample buffer and boiled at 95°C for 10 minutes. 30 μL of the sample was then loaded on 12.5% SDS-PAGE gel and run at 120 V for 70 minutes.

### Western blots

Proteins from SDS-PAGE gels were transferred to a 0.45 μm nitrocellulose membrane by wet transfer in Transfer Buffer (48 mM Tris Base, 39 mM Glycine, 0.037% SDS, 20% methanol) for 60–70 min at 10 V. Membranes were blocked overnight in a 5% (w/v) milk/TBS-T solution (5 g nonfat dry milk per 100 mL buffer, 25 mM Tris-Cl pH 8.0, 125 mM NaCl 0.1% Tween 20). The membrane was washed 3 times with TBS-T rocking for 5 min and subsequently incubated with α-FLAG-HRP primary antibody (Sigma, 1:1,000) to blot for FLAG-tagged proteins, α-VopD antibody (1:5000) (74) to blot for VopD, and anti-*E. coli* RNA polymerase β (1:3000) to blot for RNA polymerase β as loading control. All primary incubations were performed in 1X TBS-T for 1 hour rocking at room temperature. Membranes were washed 3 times with 1X TBS-T rocking for 5 min and, when necessary, incubated with goat α-mouse or goat α-rabbit IRDye 800CW secondary antibody (1:10,000, Li-COR) for 45 mins rocking at room temperature. Membranes were washed 3 times with TBS-T rocking for 5 min and then imaged on Li-COR imager.

### Yeast transformations

Wild-type *Saccharomyces cerevisiae* YHP499 cells (*MATa ura3-52 lys2-801_amber ade2-101_ochre trp1*Δ*63 his3*Δ*200 leu2*Δ*1*) (75) were grown in rich media (YPD; 1% (w/v) yeast extract, 2% (w/v) peptone, and 2% (w/v) dextrose) at 30°C overnight before back-dilution into 50 mL YPD and transformation with 100 ng plasmid DNA using the LiAc method (76). Cells from the transformation reactions were gently pelleted for 30 s at 7000 g, resuspended in 200 μL sterile water, plated on selective medium lacking uracil (Drop-out Mix Synthetic Minus Uracil; United States Biological, Salem, MA) and containing 2% (w/v) glucose, and incubated at 30°C for 3 days prior to counting colonies.

### Spot dilution assays

Spot dilution assays were performed essentially as described in (77). Briefly, *S. cerevisiae* strains were incubated overnight at 30°C with aeration in liquid media lacking uracil supplemented with 2% (w/v) raffinose. The optical density at 600 nm (OD_600_) of each culture was measured with a spectrophotometer (Eppendorf) and adjusted with sterile water to OD_600_ = 1.0 in a final volume of 200 μL. The cultures were then serially diluted 10-fold with sterile water to 10^−4^. Starting with the OD_600_ = 1.0 samples, 10 µL of each dilution was spotted onto solid media lacking uracil and supplemented with 2% (w/v) glucose or 2% (w/v) galactose. The culture spots were allowed to dry at room temperature before the plates were incubated for 2 days at 30°C. The plates were imaged using a flatbed scanner. All assays were performed in biological triplicates, and representative images are shown.

### Comparative proteomics analyses

#### Protein secretion

*V. campbellii* strains were grown for 16 hours in MLB at 30°C (LB containing 3% [w/v] NaCl). Bacterial cultures were normalized to an OD_600_ of 0.18 in 5=:□mL of MLB supplemented with 15 mM MgSO_4_ and 5 mM EGTA and incubated with constant shaking (220=:□RPM) at 30°C for 5 hours. Then, supernatant volumes equivalent to 5 OD_600_ units were filtered (0.22=:□µm), and proteins were precipitated using the deoxycholate and trichloroacetic acid method (78). The precipitated proteins were washed twice with cold acetone and then shipped to the Smoler Proteomics Center at the Technion, Israel, for analysis.

#### Proteolysis

The protein pellets were dissolved in 8.5 M Urea, 400 mM ammonium bicarbonate, and 10 mM DTT. Protein concentrations were estimated using Bradford readings. The samples were reduced (60°C for 30 minutes), modified with 35.2 mM iodoacetamide in 100 mM ammonium bicarbonate (room temperature for 30 minutes in the dark), and digested in 1.5 M Urea, 66 mM ammonium bicarbonate with modified trypsin (Promega) overnight at 37°C in a 1:50 (M/M) enzyme-to-substrate ratio. An additional trypsinization step was performed for 4 hours in a 1:100 (M/M) enzyme-to-substrate ratio. The tryptic peptides were then desalted using a homemade C18 stage tip, dried, and re-suspended in 0.1% Formic acid.

#### Mass spectrometry analysis

The resulting peptides were analyzed by LC-MS/MS using an Exploris 480 mass spectrometer (Thermo) fitted with a capillary HPLC Evosep. The peptides were loaded onto a 15 cm, ID 150 µm, 1.9-micron Endurancse column EV1137 (Evosep). The peptides were eluted with the built-in Xcalibur 15 SPD (88 min) method. Mass spectrometry was performed in a positive mode using repetitively full MS scan (m/z 350–1200) followed by High energy Collision Dissociation (HCD) of the 20 most dominant ions (>1 charges) selected from the full MS scan. A dynamic exclusion list was enabled with an exclusion duration of 30 seconds.

#### Data analysis

The mass spectrometry data were analyzed using the MaxQuant software version 2.1.1.0 for peak picking and identification using the Andromeda search engine, searching against the *V. campbellii* BB120 proteome from the NCBI-nr database, with mass tolerance of 6 ppm for the precursor masses and 20 ppm for the fragment ions. Oxidation on methionine and protein N-terminus acetylation were accepted as variable modifications, and carbamidomethyl on cysteine was accepted as static modifications. Minimal peptide length was set to seven amino acids, and a maximum of two mis-cleavages was allowed. The data were quantified by label-free analysis using MaxQuant. Peptide- and protein-level false discovery rates (FDRs) were filtered to 1% using the target-decoy strategy. The protein table was filtered to eliminate the identifications from the reverse database and common contaminants. Statistical analysis of the identification and quantization results was done using Perseus 1.6.7.0 software (79).

### T3SS Gene Homology among Vibrio species

Protein sequences from 287 publicly available complete *Vibrio* genome assemblies, previously analyzed in Simpson, *et. al.,* (80), were clustered using cd-hit (v4.8.1-2019-0228) (81) with the parameters -M 0 -g 1 -s 0.8 -c 0.65 -d 500. The dataset also included sequences encoded by the 49 T3SS genes from *Vibrio parahaemolyticus* RIMD2210633.

Sequences grouped with individual T3SS genes were designated as potential homologs. To identify more distant homologs (<65% identity), multiple sequence alignments were generated for each T3SS gene cluster using MUSCLE (v3.8.31) (82). Profile HMMs were then constructed with HMMER (v3.2.1) (83) and the complete Vibrio protein dataset was scanned using HMM search. Match scores were evaluated, and syntenic context was incorporated to confirm homolog assignments.

Sequence identity for each potential homolog was calculated by pairwise alignment against the corresponding *V. parahaemolyticus* RIMD2210633 T3SS gene using BLASTP (84, 85). For each *Vibrio* species, the highest-scoring homolog across all strains was taken as representative. The resulting homology information was summarized in a heatmap constructed through hierarchical clustering for visualization. In addition, these values were mapped onto the maximum-likelihood phylogenetic tree previously generated by Simpson, et. al., (80) and visualized using the Interactive Tree of Life - iTOL, (v5) (86). In Figure 7, the determination of whether a strain was “T3SS^+^ “ or “T3SS^−^ “ was based on the presence of T3SS gene homologs for structural genes. There was a clear shift in the number of T3SS gene homologs present in a strain, as evidenced from the table below. Thus, any strain with ≥37 predicted homologs was labeled as “T3SS^+^ “.

| Number of<br>T3SS<br>homologs | Number of<br>*Vibrio*<br>assemblies | Delineated<br>as T3SS+/- |
| --- | --- | --- |
| 2 | 3 | T3SS<sup>-</sup> |
| 5 | 3 | T3SS<sup>-</sup> |
| 6 | 8 | T3SS<sup>-</sup> |
| 7 | 117 | T3SS<sup>-</sup> |
| 8 | 3 | T3SS<sup>-</sup> |
| 9 | 26 | T3SS<sup>-</sup> |
| 10 | 13 | T3SS<sup>-</sup> |
| 11 | 6 | T3SS<sup>-</sup> |
| 12 | 8 | T3SS<sup>-</sup> |
| 37 | 2 | T3SS<sup>+</sup> |
| 38 | 3 | T3SS<sup>+</sup> |
| 39 | 1 | T3SS<sup>+</sup> |
| 40 | 1 | T3SS<sup>+</sup> |
| 41 | 1 | T3SS<sup>+</sup> |
| 42 | 7 | T3SS<sup>+</sup> |
| 43 | 27 | T3SS<sup>+</sup> |
| 44 | 6 | T3SS<sup>+</sup> |
| 45 | 3 | T3SS<sup>+</sup> |
| 46 | 14 | T3SS<sup>+</sup> |
| 47 | 12 | T3SS<sup>+</sup> |
| 48 | 3 | T3SS<sup>+</sup> |
| 49 | 20 | T3SS<sup>+</sup> |

In the comparative genomics analysis conducted we acknowledge that are a total of seven *V. parahaemolyticus* genes with less than 100% identity to the RIMD 2210633 query sequences for these genes:

|  |  |  |
| --- | --- | --- |
| VP1659 | VcrV1 | 99.67% |
| VP1662 | VcrD | 99.58% |
| VP1664 | VscX | 99.21% |
| VP1677 | VP1677 | 89.68% |
| VP1679 | VP1679 | 99.05% |
| VP1684 | VP1684 | 89.76% |
| VP1698 | ExsD | 98.19% |

This discrepancy occurred because the *V. parahaemolyticus* protein database was built from RefSeq assemblies, which include the two RIMD genomes NC_004603.1 and NC_004605.1, where locus tags follow the format VP_RSNNNNN. In contrast, the query IDs provided were in the format VPNNNN, corresponding instead to the GenBank assemblies BA000031.2 and BA000032.2 of RIMD 2210633. Consequently, the analysis compared GenBank sequences against RefSeq sequences. Any gene hits with less than full identity reflect annotation differences between GenBank and RefSeq.

### T3SS cluster comparisons

The *V. campbellii* BB120 T3SS gene cluster was identified by sequence homology to T3SS1 in *V. parahaemolyticus* RIMD 2210633 (87). Protein sequence similarity was identified using Clinker (88).

### T3SS Gene Homology among other bacterial species

Protein sequences from selected bacterial species: *Shigella flexneri* (GCF_000183785.1), *Salmonella enterica* (GCF_000006945.2), *Pseudomonas aeruginosa* (GCF_045348545.1), *Yersinia enterocolitica* (GCF_000009345.1), *Yersinia pseudotuberculosis* (GCF_000834295.1), *Yersinia pestis* (GCF_001293415.1), *Vibrio parahaemolyticus* (GCF_000196095.1), and *Vibrio campbellii* (GCF_000017705.1) — were searched for homology to the T3SS genes of *Vibrio campbellii* BB120 using the same approach described above for the *Vibrio* species.

### Identification of effector homologs using BLAST

To identify homologs of the effectors VIBHAR_05674 and VIBHAR_06684, we used the protein sequences of WP_041853546.1 and WP_012130081.1 as queries in PSI-BLAST analysis (85) until saturation (2 and 6 iterations, respectively), against the Refseq protein database (May 10, 2026) using default parameters. A maximum of 500=:□hits with a threshold of 10^−5^, a minimum 70% query coverage, and 30% identity filter were used (Datasets 3, 4).

### BMDM infection assays

#### Mice

C57BL/6 wild-type mice were bred under specific pathogen-free conditions in the animal facility at Tel Aviv University. Experiments were performed on 6- to 8-week-old female or male mice, according to the guidelines of the Institute’s Animal Ethics Committee.

#### Cell cultures

Bone marrow (BM) cells from 6- to 8-week-old mice were isolated by flushing femurs and tibias with 5 mL PBS supplemented with 0.3 (v/v) EDTA (Sartorius, 01-862-1B). The BM cells were centrifuged for 5 minutes at 400 x g and then resuspended in DMEM (Sartorius, 01-052-1A) supplemented with 10% (vol/vol) FBS and 15% L929 conditional medium (L-con). Bone marrow-derived macrophages (BMDMs) were obtained by 7 days of differentiation, as previously described (90).

#### Infection assays

*V. campbellii* strains were grown for 16 hours in MLB. Then, bacterial cultures were diluted two-fold into fresh media and incubated for an additional hour at 30°C. Approximately 3.5 × 10^4^ BMDMs were seeded into 96-well plates in triplicate in penicillin-streptomycin-free DMEM media and then infected with the indicated *V. campbellii* strains at a multiplicity of infection (MOI) ∼ 5. Plates were centrifuged for 5 minutes at 400 x g. Propidium iodide (PI; 1 μg/mL) was added to the medium 30 minutes prior to infection, and its uptake kinetics were assessed using real-time microscopy (Incucyte SX5) during incubation at 37°C. Images were acquired every 20 minutes. The addition of LPS (100 ng/mL) three hours prior to infection was used as a control for morphology changes. The data were analyzed using the Incucyte SX5 analysis software. The total BMDM cell numbers were determined using the Incucyte’s AI-driven cell segmentation analysis. Then, PI-positive (red fluorescence) cells were identified, and cell death was quantified as the percentage of PI-positive cells relative to the total cell count. The data were exported to GraphPad Prism for AUC calculations.

## Supporting information

Supporting informatio

## Data availability

RNA-seq data have been deposited at GEO under accession GSE256368. The mass spectrometry data have been deposited to the ProteomeXchange Consortium via the PRIDE (91) partner repository with the dataset identifier PXD074406. The data can be downloaded via https://ftp.pride.ebi.ac.uk/pride/data/archive/2026/03/PXD074406/.

## Acknowledgments

The authors thank Chelsea Simpson and Victoria Lydick for excellent technical support. We thank the Smoler Proteomics Center at the Technion for performing and analyzing the mass spectrometry data. Research reported in this publication was supported by: 1) the National Institute of General Medical Sciences (NIGMS) of the National Institutes of Health (NIH) under award number R35GM124698 to JVK (The content is solely the responsibility of the authors and does not necessarily represent the official views of the National Institutes of Health); 2) the US-Israel Binational Science Foundation under award number 2021733 to DS, and 3) from the Israel Science Foundation grant number 2174/22 to MG. HC was supported by a Gray Scholarship for Post-Doctoral Fellows, Gray Faculty of Medical and Health Sciences Award, Tel Aviv University; SM was supported by The Hella and Marian Gertner Fellowship and the Gray Faculty of Medical & Health Sciences Scholarship for Graduate Studies, Tel Aviv University.

## References

1. Kang H, Yu Y, Liao M, Wang Y, Yang G, Zhang Z, et al. Physiology, metabolism, antibiotic resistance, and genetic diversity of Harveyi clade bacteria isolated from coastal mariculture system in China in the last two decades. Frontiers in Marine Science. 2022;9:932255-.

2. Kumar S, Kumar CB, Rajendran V, Abishaw N, Anand PSS, Kannapan S, et al. Delineating virulence of Vibrio campbellii: a predominant luminescent bacterial pathogen in Indian shrimp hatcheries. Scientific Reports 2021;11(1):1–16.

3. Liu J, Zhao Z, Deng Y, Shi Y, Liu Y, Wu C, et al. Complete genome sequence of Vibrio campbellii LMB 29 isolated from red drum with four native megaplasmids. Frontiers in Microbiology. 2017;8(OCT):290197-.

4. Srisangthong I, Sangseedum C, Chaichanit N, Surachat K, Suanyuk N, Mittraparp-arthorn P. Characterization and Genome Analysis of Vibrio campbellii Lytic Bacteriophage OPA17. Microbiology Spectrum. 2023;11(2).

5. Zhang XH, He X, Austin B. Vibrio harveyi: a serious pathogen of fish and invertebrates in mariculture. Marine Life Science & Technology. 2020;2(3):231-.

6. Bachand PT, Tallman JJ, Powers NC, Woods M, Azadani DN, Zimba PV, et al. Genomic identification and characterization of co-occurring Harveyi clade species following a vibriosis outbreak in Pacific white shrimp, Penaeus (litopenaeus) vannamei. Aquaculture. 2020;518:734628-.

7. Urbanczyk H, Ogura Y, Hayashi T. Taxonomic revision of Harveyi clade bacteria (family Vibrionaceae) based on analysis of whole genome sequences. International Journal of Systematic and Evolutionary Microbiology. 2013;63(PART7):2742–51.

8. Coburn B, Sekirov I, Finlay BB. Type III Secretion Systems and Disease. Clinical Microbiology Reviews. 2007;20(4):535-.

9. Puhar A, Sansonetti PJ. Type III secretion system. Current Biology. 2014;24(17):R784–R91.

10. Trosky JE, Liverman ADB, Orth K. Yersinia outer proteins: Yops. Cellular Microbiology. 2008;10(3):557–65.

11. Letchumanan V, Chan KG, Lee LH. Vibrio parahaemolyticus: a review on the pathogenesis, prevalence, and advance molecular identification techniques. Front. Microbiol. 5:705. doi: 10.3389/fmicb.2014.00705

12. Lee VT, Smith RS, Tümmler B, Lory S. Activities of Pseudomonas aeruginosa Effectors Secreted by the Type III Secretion System In Vitro and during Infection. INFECTION AND IMMUNITY. 2005;73(3):1695–705.

13. De Nisco NJ, Kanchwala M, Li P, Fernandez J, Xing C, Orth K. Cytotoxic Vibrio T3SS1 Rewires Host Gene Expression to Subvert Cell Death Signaling and Activate Cell Survival Networks. Science signaling. 2017;10(479).

14. Agbor TA, McCormick BA. Salmonella Effectors: Important players modulating host cell function during infection. Cellular Microbiology. 2011;13(12):1858-.

15. Iriarte M, Cornelis GR. YopT, a new Yersinia Yop effector protein, affects the cytoskeleton of host cells. Molecular Microbiology. 1998;29(3):915–29.

16. Makino K, Oshima K, Kurokawa K, Yokoyama K, Uda T, Tagomori K, et al. Genome sequence of Vibrio parahaemolyticus: a pathogenic mechanism distinct from that of V cholerae. Lancet. 2003;361(9359):743–9.

17. Broberg CA, Calder TJ, Orth K. Vibrio parahaemolyticus cell biology and pathogenicity determinants. Microbes Infect. 2011;13(12-13):992–1001.

18. Ritchie JM, Rui H, Zhou X, Iida T, Kodoma T, Ito S, et al. Inflammation and disintegration of intestinal villi in an experimental model for Vibrio parahaemolyticus-induced diarrhea. PLoS pathogens. 2012;8(3):e1002593.

19. Zhang L, Krachler AM, Broberg CA, Li Y, Mirzaei H, Gilpin CJ, et al. Type III effector VopC mediates invasion for Vibrio species. Cell Rep. 2012;1(5):453–60.

20. Yang H, de Souza Santos M, Lee J, Law HT, Chimalapati S, Verdu EF, et al. A Novel Mouse Model of Enteric Vibrio parahaemolyticus Infection Reveals that the Type III Secretion System 2 Effector VopC Plays a Key Role in Tissue Invasion and Gastroenteritis. mBio. 2019;10(6).

21. Zhang L, Orth K. Virulence determinants for Vibrio parahaemolyticus infection. Current Opinion in Microbiology. 2013;16(1):70–7.

22. Dong X, Wang H, Zou P, Chen J, Liu Z, Wang X, et al. Complete genome sequence of Vibrio campbellii strain 20130629003S01 isolated from shrimp with acute hepatopancreatic necrosis disease. Gut Pathogens. 2017;9(1):1–5.

23. Liang J, Liu J, Wang X, Sun H, Zhang Y, Ju F, et al. Genomic Analysis Reveals Adaptation of Vibrio campbellii to the Hadal Ocean. Appl Environ Microbiol 88:e00575–22. 10.1128/aem.00575-22

24. Lin B, Wang Z, Malanoski AP, O’Grady EA, Wimpee CF, Vuddhakul V, et al. Comparative genomic analyses identify the Vibrio harveyi genome sequenced strains BAA-1116 and HY01 as Vibrio campbellii. Environmental Microbiology Reports. 2010;2(1):81-.

25. Darshanee Ruwandeepika HA, Karunasagar I, Bossier P, Defoirdt T. Expression and Quorum Sensing Regulation of Type III Secretion System Genes of Vibrio harveyi during Infection of Gnotobiotic Brine Shrimp. PloS one. 2015;10(12).

26. Yarbrough ML, Li Y, Kinch LN, Grishin NV, Ball HL, Orth K. AMPylation of Rho GTPases by Vibrio VopS disrupts effector binding and downstream signaling. Science. 2009;323(5911):269–72.

27. Sreelatha A, Bennett TL, Carpinone EM, O’Brien KM, Jordan KD, Burdette DL, et al. Vibrio effector protein VopQ inhibits fusion of V-ATPase-containing membranes. Proceedings of the National Academy of Sciences of the United States of America. 2015;112(1):100–5.

28. Salomon D, Guo Y, Kinch LN, Grishin NV, Gardner KH, Orth K. Effectors of animal and plant pathogens use a common domain to bind host phosphoinositides. Nature Communications 2013 4:1. 2013;4(1):1–10.

29. Broberg CA, Zhang L, Gonzalez H, Laskowski-Arce MA, Orth K. A Vibrio effector protein is an inositol phosphatase and disrupts host cell membrane integrity. Science. 2010;329(5999):1660–2.

30. Waters CM, Wu JT, Ramsey ME, Harris RC, Bassler BL. Control of the Type 3 Secretion System in Vibrio harveyi by Quorum Sensing through Repression of ExsA. Applied and Environmental Microbiology. 2010;76(15):4996-.

31. Henke JM, Bassler BL. Quorum sensing regulates type III secretion in Vibrio harveyi and Vibrio parahaemolyticus. Journal of bacteriology. 2004;186(12):3794–805.

32. Paul P, Podicheti R, Geyman LJ, Baker EN, Papenfort K, Rusch DB, et al. Quorum sensing employs a dual regulatory mechanism to repress T3SS gene expression. mBio. 2025.

33. Van Kessel JC, Rutherford ST, Shao Y, Utria AF, Bassler BL. Individual and Combined Roles of the Master Regulators AphA and LuxR in Control of the Vibrio harveyi Quorum-Sensing Regulon. Journal of Bacteriology. 2013;195(3):436-.

34. Dewoody RS, Merritt PM, Marketon MM. Regulation of the Yersinia type III secretion system: Traffic control. Frontiers in Cellular and Infection Microbiology. 2013;4(FEB):36455-.

35. Hotinger JA, Pendergrass HA, May AE. Molecular Targets and Strategies for Inhibition of the Bacterial Type III Secretion System (T3SS); Inhibitors Directly Binding to T3SS Components. Biomolecules 2021, Vol 11, Page 316. 2021;11(2):316-.

36. Wimmi S, Balinovic A, Jeckel H, Selinger L, Lampaki D, Eisemann E, et al. Dynamic relocalization of cytosolic type III secretion system components prevents premature protein secretion at low external pH. Nature Communications 2021 12:1. 2021;12(1):1–14.

37. Diepold A, Armitage JP. Type III secretion systems: the bacterial flagellum and the injectisome. Philosophical Transactions of the Royal Society B: Biological Sciences. 2015;370(1679).

38. Deng W, Marshall NC, Rowland JL, McCoy JM, Worrall LJ, Santos AS, et al. Assembly, structure, function and regulation of type III secretion systems. Nature Reviews Microbiology 2017 15:6. 2017;15(6):323–37.

39. McMackin EAW, Djapgne L, Corley JM, Yahr TL. Fitting Pieces into the Puzzle of Pseudomonas aeruginosa Type III Secretion System Gene Expression. 2019.

40. Schwiesow L, Lam H, Dersch P, Auerbuch V. Yersinia Type III Secretion System Master Regulator LcrF. Journal of Bacteriology. 2016;198(4):604-.

41. Liu AC, Thomas NA. Transcriptional profiling of Vibrio parahaemolyticus exsA reveals a complex activation network for type III secretion. Frontiers in Microbiology. 2015;6(OCT):1089-.

42. Kodama T, Yamazaki C, Park KS, Akeda Y, Iida T, Honda T. Transcription of Vibrio parahaemolyticus T3SS1 genes is regulated by a dual regulation system consisting of the ExsACDE regulatory cascade and H-NS. FEMS Microbiology Letters. 2010;311(1):10–7.

43. Okada N, Iida T, Park KS, Goto N, Yasunaga T, Hiyoshi H, et al. Identification and characterization of a novel type III secretion system in trh-positive Vibrio parahaemolyticus strain TH3996 reveal genetic lineage and diversity of pathogenic machinery beyond the species level. Infection and immunity. 2009;77(2):904–13.

44. Zhang Y, Deng Y, Feng J, Guo Z, Chen H, Wang B, et al. Functional characterization of VscCD, an important component of the type Ⅲ secretion system of Vibrio harveyi. Microb Pathog. 2021;157:104965.

45. Sarty D, Baker NT, Thomson EL, Rafuse C, Ebanks RO, Graham LL, et al. Characterization of the type III secretion associated low calcium response genes of Vibrio parahaemolyticus RIMD2210633. Can J Microbiol. 2012;58(11):1306–15.

46. Barve SS, Straley SC. lcrR, a low-Ca2(+)-response locus with dual Ca2(+)-dependent functions in Yersinia pestis. Journal of Bacteriology. 1990;172(8):4661-.

47. Dasgupta N, Ashare A, Hunninghake GW, Yahr TL. Transcriptional induction of the Pseudomonas aeruginosa type III secretion system by low Ca2+ and host cell contact proceeds through two distinct signaling pathways. Infection and Immunity. 2006;74(6):3334–41.

48. Liu J, Lu SY, Orfe LH, Ren CH, Hu CQ, Call DR, et al. ExsE is a negative regulator for T3SS gene expression in Vibrio alginolyticus. Frontiers in Cellular and Infection Microbiology. 2016;6(DEC):177-.

49. Barbieri JT, Sun J. Pseudomonas aeruginosa ExoS and ExoT. Reviews of Physiology, Biochemistry and Pharmacology. 2004;152:79–92.

50. Ganesan AK, Frank DW, Misra RP, Schmidt G, Barbieri JT. Pseudomonas aeruginosa Exoenzyme S ADP-ribosylates Ras at Multiple Sites. Journal of Biological Chemistry. 1998;273(13):7332–7.

51. Sundin C, Hallberg B, Forsberg Ãk. ADP-ribosylation by exoenzyme T of Pseudomonas aeruginosa induces an irreversible effect on the host cell cytoskeleton in vivo. FEMS Microbiology Letters. 2004;234(1):87–91.

52. Wang J, Chitsaz F, Derbyshire MK, Gonzales NR, Gwadz M, Lu S, et al. The conserved domain database in 2023. Nucleic Acids Research. 2023/01/06;51(D1).

53. Lenstra Tineke L, Coulon A, Chow Carson C, Larson Daniel R. Single-Molecule Imaging Reveals a Switch between Spurious and Functional ncRNA Transcription. Molecular Cell. 2015;60(4).

54. Higa N, Toma C, Koizumi Y, Nakasone N, Nohara T, Masumoto J, et al. Vibrio parahaemolyticus Effector Proteins Suppress Inflammasome Activation by Interfering with Host Autophagy Signaling. PLoS Pathogens. 2013 Jan 24;9(1).

55. Hoffmann M, Monday SR, Allard MW, Strain EA, Whittaker P, Naum M, et al. Vibrio caribbeanicus sp. nov., isolated from the marine sponge Scleritoderma cyanea. International Journal of Systematic and Evolutionary Microbiology. 2012/08/01;62(Pt_8).

56. Huang B, Feng Y, Qin Z, Yu Z, Tian Z, Peng K, et al. Vibrio misgurnus sp. nov., a new pathogen of cultured loach (Misgurnus anguillicaudatus), closely related to Vibrio cholerae. Aquaculture. 2024/05/30;586.

57. Budiyansah H, Mursalim MF, Chokmangmeepisarn P, Rung-ruangkijkrai T, Mabrok M, Khan MIR, et al. Unveiling the threat: Aeromonas schubertii, an emerging bacterial pathogen inducing white nodule lesions in snakehead fish (Channa striata) at nursery farm in Thailand. Aquaculture International 2025 33:6. 2025-08-19;33(6).

58. Chen F, Sun J, Han Z, Yang X, Xian J-a, Lv A, et al. Frontiers | Isolation, Identification and Characteristics of Aeromonas veronii From Diseased Crucian Carp (Carassius auratus gibelio). Frontiers in Microbiology. 2019/11/26;10.

59. Hawke JP, Kent M, Rogge M, Baumgartner W, Wiles J, Shelley J, et al. Edwardsiellosis Caused by Edwardsiella ictaluri in Laboratory Populations of Zebrafish Danio rerio. Journal of aquatic animal health. 2013 Sep;25(3).

60. Kumar S, Kumar CB, Rajendran V, Abishaw N, Anand PSS, Kannapan S, et al. Delineating virulence of Vibrio campbellii: a predominant luminescent bacterial pathogen in Indian shrimp hatcheries. Sci Rep. 2021;11(1):15831.

61. Ono T, Park KS, Ueta M, Iida T, Honda T. Identification of proteins secreted via Vibrio parahaemolyticus type III secretion system 1. Infection and immunity. 2006;74(2):1032–42.

62. Pan X, Yang Y, Zhang JR. Molecular basis of host specificity in human pathogenic bacteria. Emerg Microbes Infect. 2014;3(3):e23.

63. Letchumanan V, Chan K-G, Lee L-H. Vibrio parahaemolyticus: a review on the pathogenesis, prevalence, and advance molecular identification techniques. Frontiers in microbiology. 2014;Volume 5 - 2014.

64. Letchumanan V, Chan KG, Lee LH. Vibrio parahaemolyticus: a review on the pathogenesis, prevalence, and advance molecular identification techniques. Frontiers in microbiology. 2014;5:705.

65. Restrepo-Benavides M, Lozano-Arce D, Gonzalez-Garcia LN, Báez-Aguirre F, Ariza-Aranguren G, Faccini D, et al. Unveiling potential virulence determinants in *Vibrio* isolates from *Anadara tuberculosa* through whole genome analyses. Microbiology Spectrum. 2024;12(2):e02928–23.

66. Baker-Austin C, Oliver JD, Alam M, Ali A, Waldor MK, Qadri F, et al. Vibrio spp. infections. Nature Reviews Disease Primers. 2018;4(1):1–19.

67. Lin B, Wang Z, Malanoski AP, O’Grady EA, Wimpee CF, Vuddhakul V, et al. Comparative genomic analyses identify the Vibrio harveyi genome sequenced strains BAA-1116 and HY01 as Vibrio campbellii. Environ Microbiol Rep. 2010;2(1):81–9.

68. Chaparian RR, Olney SG, Hustmyer CM, Rowe-Magnus DA, van Kessel JC. Integration host factor and LuxR synergistically bind DNA to coactivate quorum-sensing genes in Vibrio harveyi. Mol Microbiol. 2016;101(5):823–40.

69. Rutherford ST, van Kessel JC, Shao Y, Bassler BL. AphA and LuxR/HapR reciprocally control quorum sensing in vibrios. Genes Dev. 2011;25(4):397–408.

70. Bolger AM, Lohse M, Usadel B. Trimmomatic: a flexible trimmer for Illumina sequence data. Bioinformatics. 2014/08/01;30(15).

71. Langmead B, Salzberg SL, Langmead B, Salzberg SL. Fast gapped-read alignment with Bowtie 2. Nature Methods 2012 9:4. 2012-03-04;9(4).

72. Liao Y, Smyth GK, Shi W. featureCounts: an efficient general purpose program for assigning sequence reads to genomic features. Bioinformatics. 2014/04/01;30(7).

73. Love MI, Huber W, Anders S. Moderated estimation of fold change and dispersion for RNA-seq data with DESeq2. Genome Biology. 2014 Dec 5;15(12).

74. Henke JM, Bassler BL. Quorum sensing regulates type III secretion in Vibrio harveyi and Vibrio parahaemolyticus. J Bacteriol. 2004;186(12):3794–805.

75. Sikorski RS, Hieter P. A system of shuttle vectors and yeast host strains designed for efficient manipulation of DNA in Saccharomyces cerevisiae. Genetics. 1989;122(1):19–27.

76. Gietz RD, Schiestl RH, Willems AR, Woods RA. Studies on the transformation of intact yeast cells by the LiAc/SS-DNA/PEG procedure. Yeast (Chichester, England). 1995;11(4):355–60.

77. Nickens DG, Bochman ML. Genetic and biochemical interactions of yeast DNA helicases. Methods (San Diego, Calif). 2022;204:234–40.

78. Bensadoun A, Weinstein D. Assay of proteins in the presence of interfering materials. Analytical Biochemistry. 1976/01/01;70(1).

79. Tyanova S, Temu T, Sinitcyn P, Carlson A, Hein MY, Geiger T, et al. The Perseus computational platform for comprehensive analysis of (prote)omics data. Nature Methods 2016 13:9. 2016-06-27;13(9).

80. Simpson CA, Petersen BD, Haas NW, Geyman LJ, Lee AH, Podicheti R, et al. The quorum-sensing systems of Vibrio campbellii DS40M4 and BB120 are genetically and functionally distinct. Environmental microbiology. 2021 Jun 7;23(9).

81. Li W, Godzik A. Cd-hit: a fast program for clustering and comparing large sets of protein or nucleotide sequences. Bioinformatics. 2006/07/01;22(13).

82. Edgar RC. MUSCLE: multiple sequence alignment with high accuracy and high throughput. Nucleic Acids Research. 2004/03/01;32(5).

83. Eddy SR. Accelerated Profile HMM Searches. PLOS Computational Biology. Oct 20, 2011;7(10).

84. Altschul SF, Gish W, Miller W, Myers EW, Lipman DJ. Basic local alignment search tool. Journal of Molecular Biology. 1990/10/05;215(3).

85. Altschul SF, Madden TL, Schäffer AA, Zhang J, Zhang Z, Miller W, et al. Gapped BLAST and PSI-BLAST: a new generation of protein database search programs. Nucleic Acids Research. 1997 Sep 1;25(17).

86. Letunic I, Bork P. Interactive Tree Of Life (iTOL) v5: an online tool for phylogenetic tree display and annotation. Nucleic Acids Research. 2021/07/02;49(W1).

87. Ono T, Park K-S, Ueta M, Iida T, Honda T. Identification of Proteins Secreted via Vibrio parahaemolyticus Type III Secretion System 1. Infection and Immunity. 2006 Feb;74(2).

88. Belt Mvd, Gilchrist C, Booth TJ, Chooi Y-H, Medema MH, Alanjary M. CAGECAT: The CompArative GEne Cluster Analysis Toolbox for rapid search and visualisation of homologous gene clusters. BMC Bioinformatics. 2023 May 3;24(1).

89. Altschul SF, Gish W, Miller W, Myers EW, Lipman DJ. Basic local alignment search tool. J Mol Biol. 1990;215(3):403–10.

90. Erlich Z, Shlomovitz I, Edry-Botzer L, Cohen H, Frank D, Wang H, et al. Macrophages, rather than DCs, are responsible for inflammasome activity in the GM-CSF BMDC model. Nature Immunology 2019 20:4. 2019-02-11;20(4).

91. Perez-Riverol Y, Bandla C, Kundu DJ, Kamatchinathan S, Bai J, Hewapathirana S, et al. The PRIDE database at 20 years: 2025 update. Nucleic Acids Res. 2025;53(D1):D543–D53.

