## Supporting informatio for "*Vibrio campbellii* encodes a distinct set of type III secretion system effectors that mediate cytotoxicity in eukaryotic host models"

**Supplemental Figures S1-S6**

**Datasets S1-S4**

**Tables S1-S2**

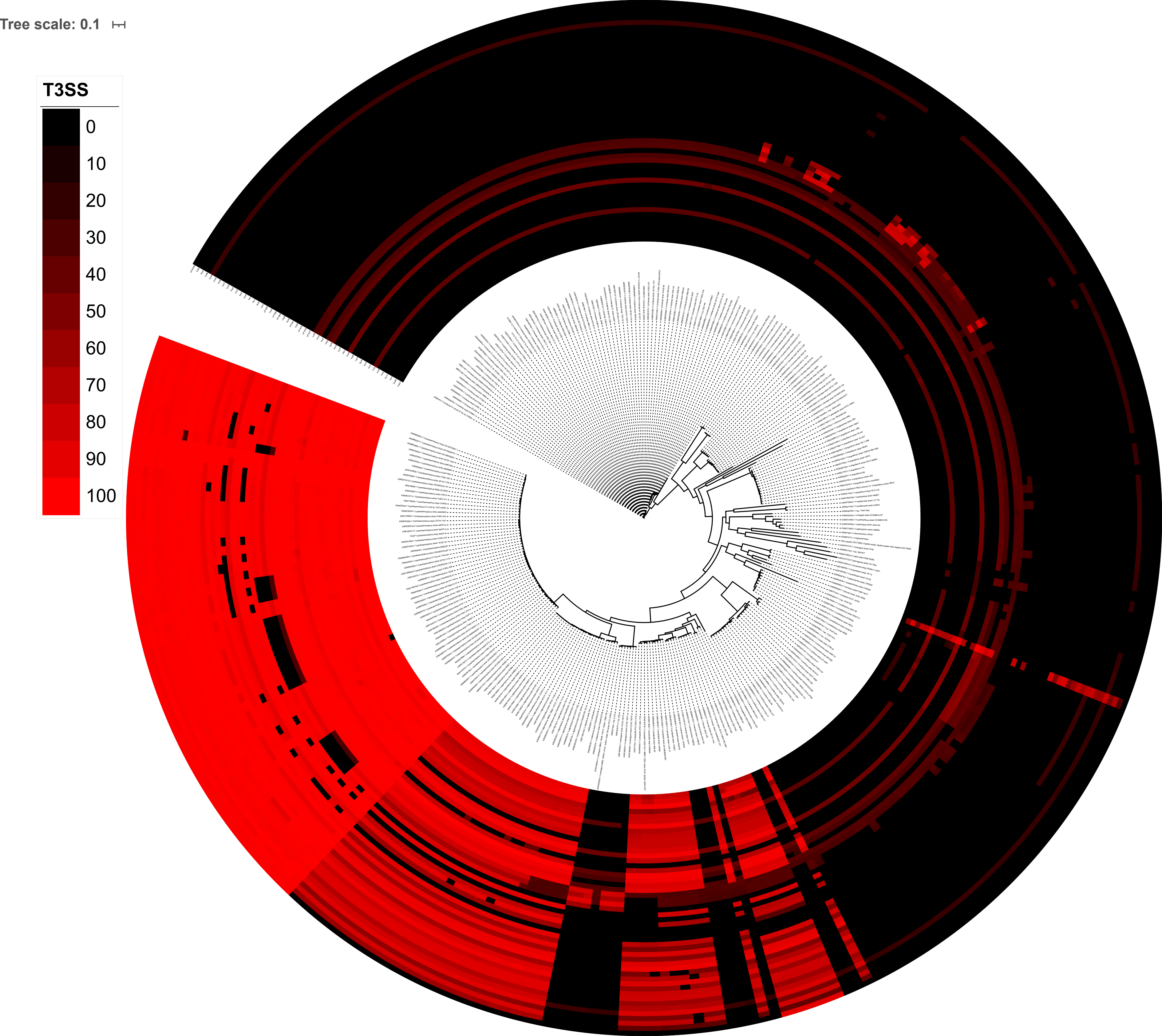
Tree scale: 0.1
T3SS
0
10
20
30
40
50
60
70
80
90
100

**Figure S1.** Comparative genomics analysis shows the T3SS1 genes conservation in the family *Vibrionaceae*. *Vibrio parahaemolyticus* RIMD2210633 has been used as the reference strain for the comparative genomics analysis.

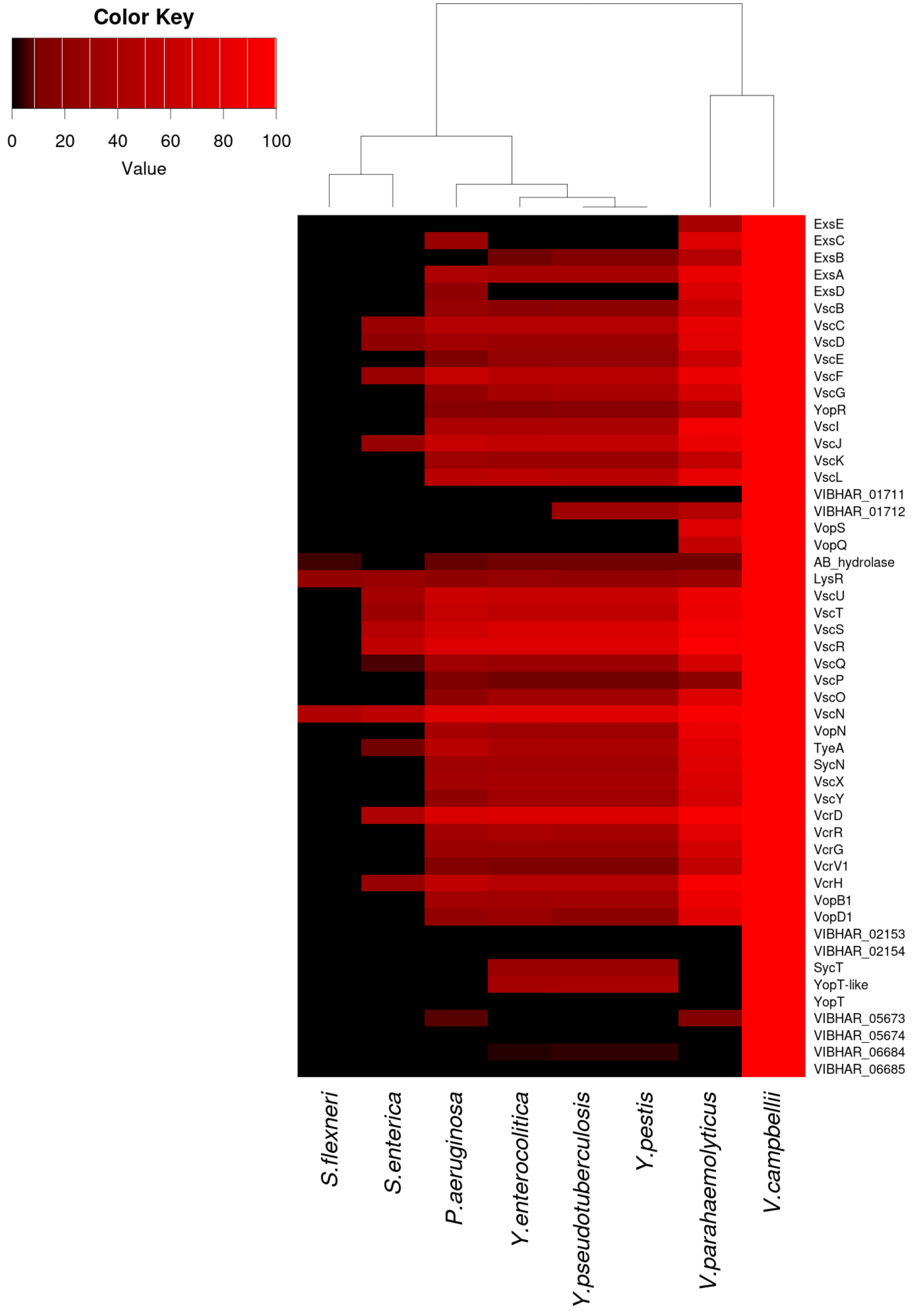


**Figure S2. Comparative genomics analysis shows conservation of the T3SS genes.** The percent amino acid identity for each T3SS gene represents the highest scoring homolog across all strains for that species compared to the reference strain *V. campbellii* BB120. The dendrogram represents the phylogenetic relationship among the bacterial species.

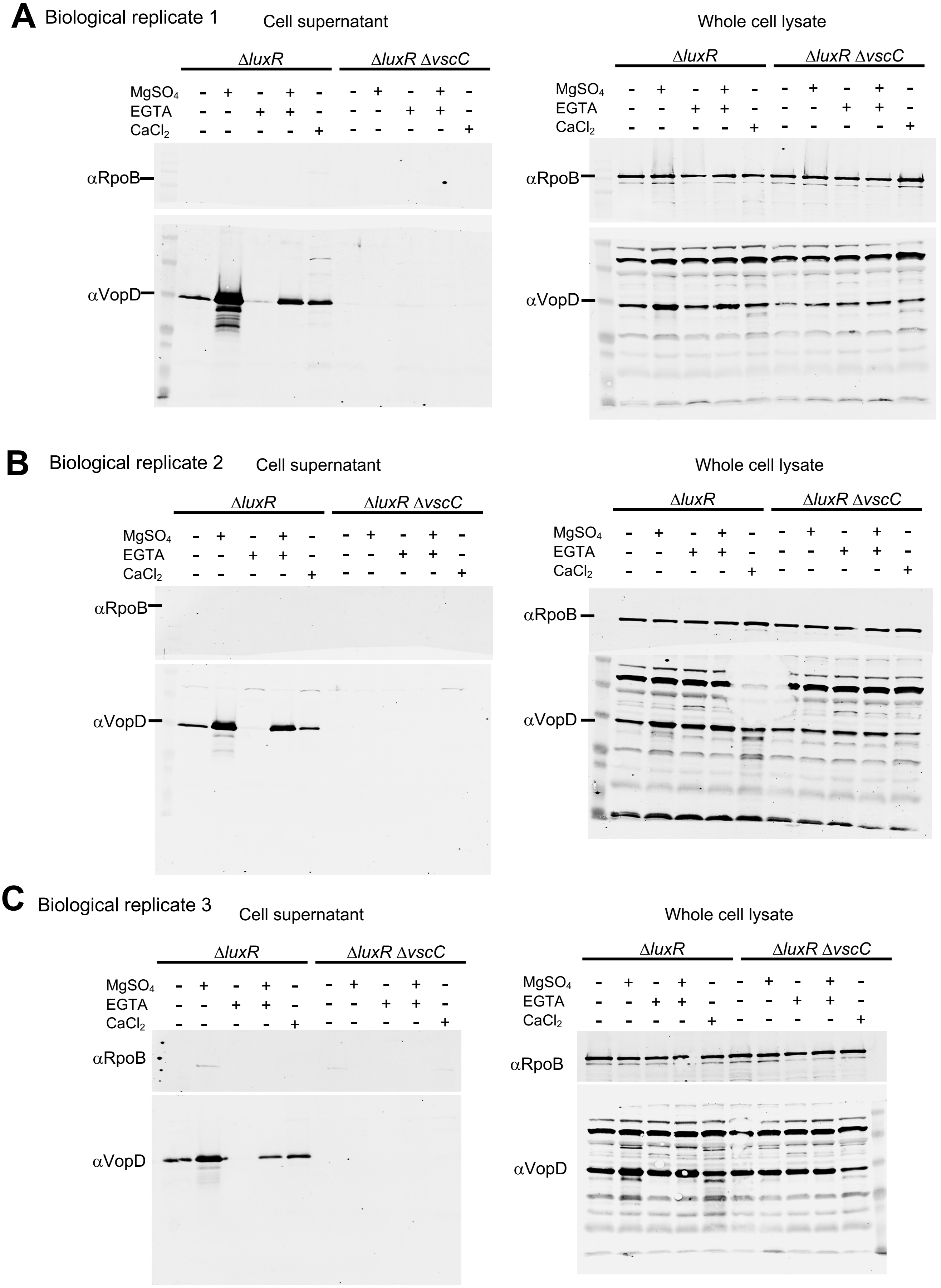


**Figure S3.** Secretion assays followed by Western Blot on cellular supernatants and whole cell lysates in Δ*luxR* and Δ*luxR* Δ*vscC* strains depict VopD secretion under different media conditions including LM only, LM+15 mM $MgSO_4$, LM+5 mM EGTA, LM+15 mM $MgSO_4$+5 mM EGTA, and LM+15 mM $CaCl_2$. VopD was blotted with anti-VopD antibody. RNA polymerase β subunit (Rpoβ) was used as loading control and blotted with anti-Rpoβ antibody.

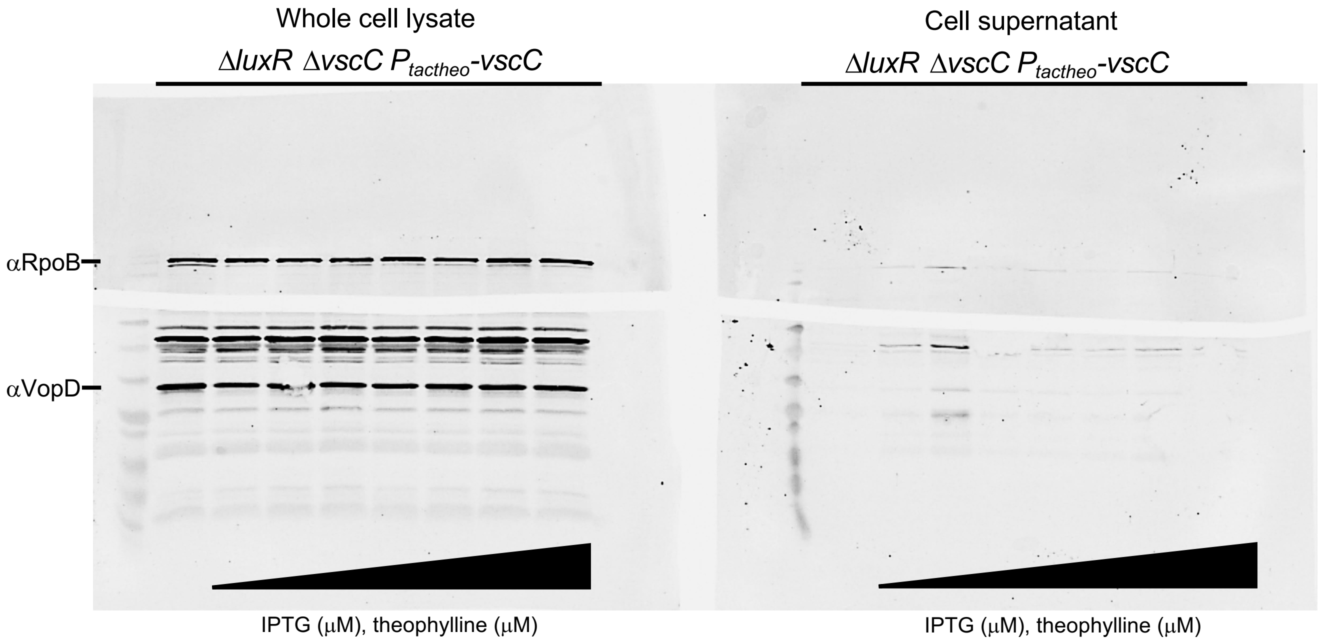


**Figure S4.** Secretion assays followed by western blot on cellular supernatants and whole cell lysates in Δ*luxR* Δ*vscC* strain with *vscC* complemented ectopically from an IPTG and theophylline induced promoter ($P_{tactheo}$-*vscC,* pPP99) depict VopD secretion in media LM+15 mM $MgSO_4$. Ectopic transcription and translation of *vscC* was titrated sequentially using 10 μM, or 100 μM, or 1000 μM IPTG in combination with 10 μM, or 100 μM, or 1000 μM theophylline. VopD was blotted with anti-VopD antibody. RNA polymerase β subunit (Rpoβ) was used as loading control and blotted with anti-Rpoβ antibody. One representative image of triplicate biological experiments is shown.

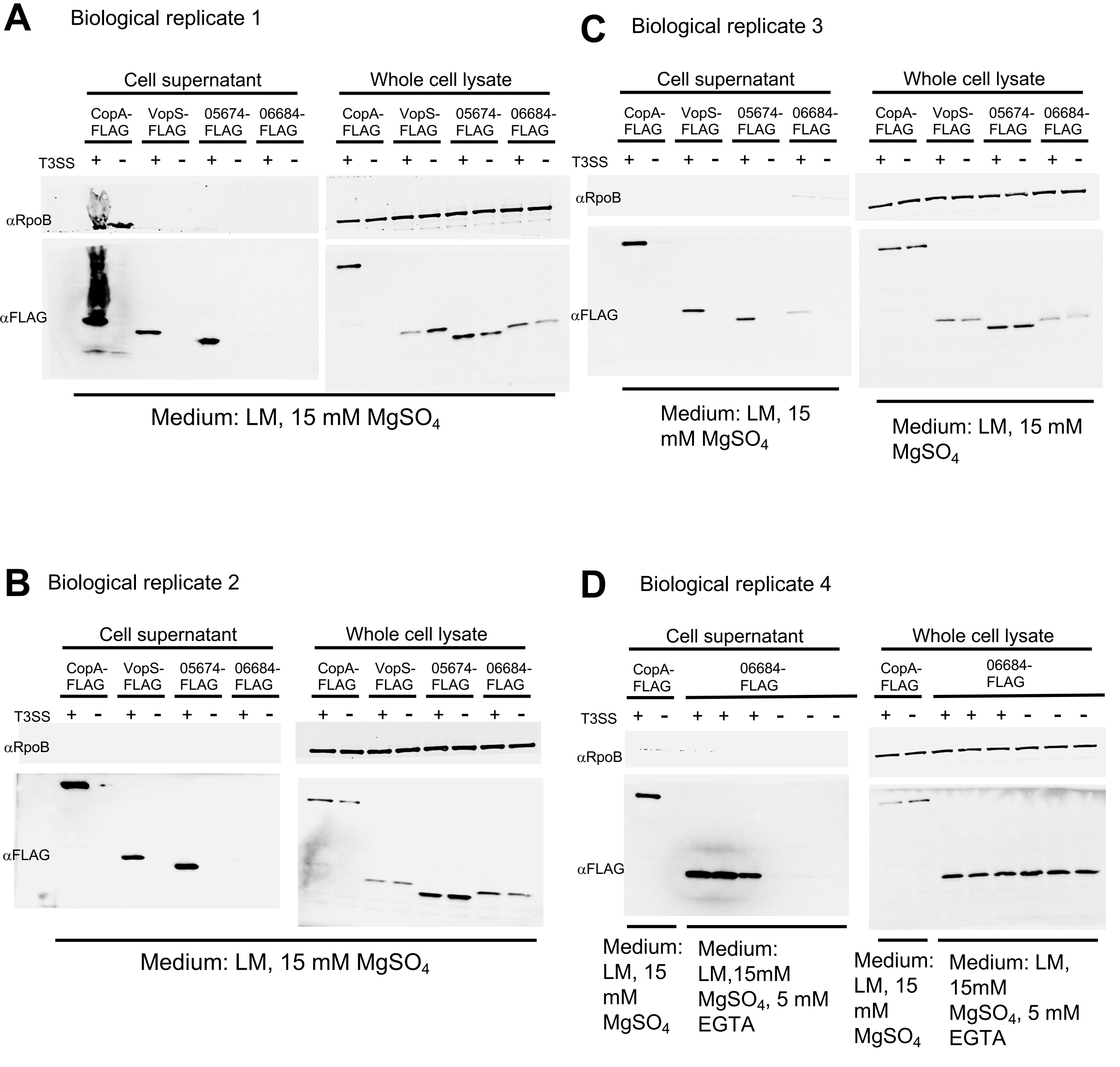


**Figure S5.** Secretion assay followed by western blot with T3SS$^+$ ($\Delta$*luxR*) and T3SS$^-$ ($\Delta$*luxR* $\Delta$*vscC*) strains on the whole cell lysate and cell supernatant. The putative effector proteins were FLAG-tagged in the C-terminus and overexpressed in an ectopic plasmid. $P_{tactheo}$-*copA-FLAG* (pPP67) was induced with 100 μM IPTG and 10 μM theophylline, $P_{tactheo}$-*vopS-FLAG* (pPP68), $P_{tactheo}$-*VIBHAR_05674-FLAG* (pPP71) were induced with 10 μM IPTG and 1000 μM theophylline, and $P_{tactheo}$-*VIBHAR_06684-FLAG* (pPP72) was induced with 100 μM IPTG and 1000 μM theophylline FLAG-tagged proteins were blotted with anti-FLAG antibody. RNA polymerase β subunit (Rpoβ) was used as loading control and blotted with anti-Rpoβ antibody.

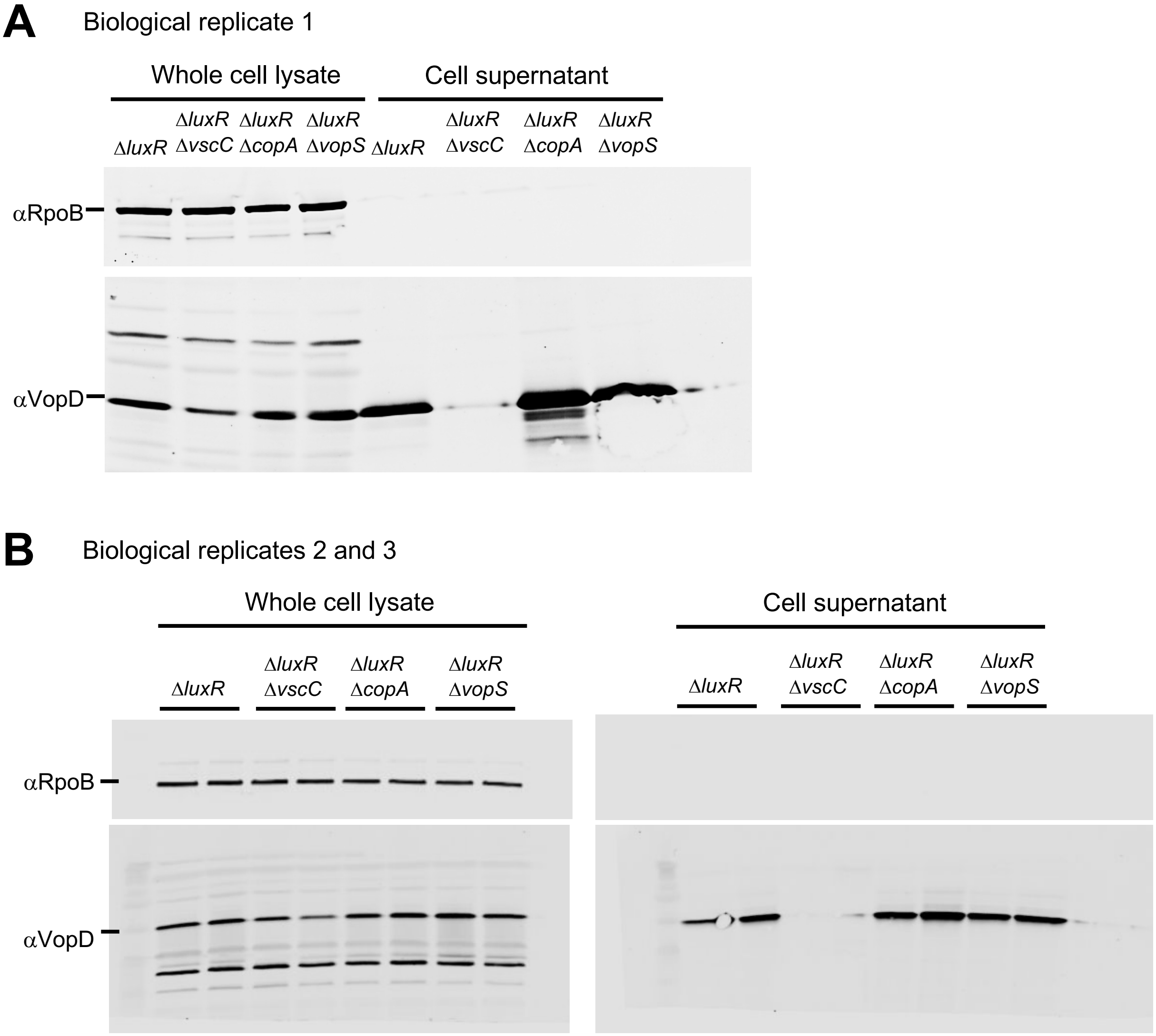


**Figure S6.** Secretion assays followed by Western Blot on cellular supernatants and whole cell lysates of *ΔluxR, ΔluxR ΔvscC, ΔluxR ΔcopA,* and *ΔluxR ΔvopS* strains depict VopD secretion in media LM+15 mM MgSO4. VopD was blotted with α-VopD antibody. RNA polymerase β subunit (Rpoβ) was used as loading control and blotted with α-Rpoβ antibody.

**Dataset S1.** RNA-seq differential gene expression analysis comparing *V. campbellii* Δ*luxR* versus Δ*luxR* Δ*vscC* strains.

**Dataset S2.** Protein table results of LC-MS/MS analysis of secreted proteins extracted from *V. campbellii* Δ*luxR* versus Δ*luxR* Δ*vscC* strain supernatants.

**Dataset S3.** PSI-BLAST analysis hits table results to identify homologs of VIBHAR_06684 (WP_012130081).

**Dataset S4.** PSI-BLAST analysis hits table results to identify homologs of VIBHAR_05674 (WP_041853546).

**Table S1.** *V. campbellii* BB120 strains used in this study.

| | Genotype | Reference |
|---|---|---|
| BB120 | wildtype | (1,2,3) |
| KM669 | Δ*luxR* | (1,2,3) |
| PPVh128 | Δ*exsA* | (4) |
| PPVh129 | Δ*luxR* Δ*exsA* | (4) |
| PPVh251 | Wild-type with chromosomal deletion of *vscC* locus | This study |
| PPVh252 | Δ*luxR* with chromosomal deletion of *vscC* locus | This study |
| PPVh319 | Δ*luxR* with ectopically expressed C-terminal FLAG tagged *copA* under $P_{tac\text{-}theo}$ | This study |
| PPVh321 | Δ*luxR* Δ*vscC* with ectopically expressed C-terminal FLAG tagged *copA* under $P_{tac\text{-}theo}$ | This study |
| PPVh322 | Δ*luxR* with ectopically expressed C-terminal FLAG tagged *vopS* under $P_{tac\text{-}theo}$ | This study |
| PPVh324 | Δ*luxR* Δ*vscC* with ectopically expressed C-terminal FLAG tagged *vopS* under $P_{tac\text{-}theo}$ | This study |
| PPVh328 | Δ*luxR* with ectopically expressed C-terminal FLAG tagged VIBHAR_05674 under $P_{tac\text{-}theo}$ | This study |
| PPVh330 | Δ*luxR* Δ*vscC* with ectopically expressed C-terminal FLAG tagged VIBHAR_05674 under $P_{tac\text{-}theo}$ | This study |
| PPVh331 | Δ*luxR* with ectopically expressed C-terminal FLAG tagged VIBHAR_06684 under $P_{tac\text{-}theo}$ | This study |
| PPVh333 | Δ*luxR* Δ*vscC* with ectopically expressed C-terminal FLAG tagged VIBHAR_06684 under $P_{tac\text{-}theo}$ | This study |
| PPVh410 | *VIBHAR_05674* deleted in WT BB120 background | This study |
| PPVh412 | *luxR* deleted in *VIBHAR_05674* deletion background | This study |
| PPVh414 | *VIBHAR_06684* deleted in a *luxR* deletion background | This study |
| PPVh424 | *copA* (VIBHAR_01711) deleted in the *luxR* deletion background | This study |
| PPVh426 | *vopS* (VIBHAR_01713) deleted in the *luxR* deletion background | This study |

**Table S2.** Plasmids used in this study.

| Strain | Genotype | Reference |
|---|---|---|
| pRE112 | gene deletion vector, $cm^R$ (chloramphenicol resistant), *sacB* (sucrose intolerant), R6Kg origin | (5, 6) |
| pFOG | gene deletion vector, $gent^R$ (gentamycin resistant), *sacB* (sucrose intolerant), anhydrous tetracycline induced expression *I-sceI* (double stranded DNA breaks at I-SceI recognition sequence), R6Kg origin | (7) |
| pPP67 | C-terminal FLAG tagged *copA* expressed under $P_{tac\text{-}theo}$ in pMMB67EH-$gent^R$ backbone | This study and (8) |
| pPP68 | C-terminal FLAG tagged *vopS* expressed under $P_{tac\text{-}theo}$ in pMMB67EH-$gent^R$ backbone | This study and (8) |
| pPP71 | C-terminal FLAG tagged VIBHAR_05674 expressed under $P_{tac\text{-}theo}$ in pMMB67EH-$gent^R$ backbone | This study and (8) |
| pPP72 | C-terminal FLAG tagged VIBHAR_06684 expressed under $P_{tac\text{-}theo}$ in pMMB67EH-$gent^R$ backbone | This study and (8) |
| pPP75 | *VIBHAR_05674* deletion construct in pRE112, $Cm^R$ | This study |
| pPP76 | *VIBHAR_06684* deletion construct in pRE112, $Cm^R$ | This study |
| pRC50 | *luxR* deletion construct in pRE112, $Cm^R$ | (5, 6) |
| pPP95 | *copA* deletion construct in pFOG, $Gentamycin^R$ | This study |
| pPP96 | *vopS* deletion construct in pFOG, $Gentamycin^R$ | This study |

### References


1. Henke JM, Bassler BL. 2004. Quorum sensing regulates type III secretion in Vibrio harveyi and Vibrio parahaemolyticus. J Bacteriol 186:3794-805.
2. van Kessel JC, Rutherford ST, Shao Y, Utria AF, Bassler BL. 2013. Individual and combined roles of the master regulators AphA and LuxR in control of the Vibrio harveyi quorum-sensing regulon. J Bacteriol 195:436-43.
3. Waters CM, Wu JT, Ramsey ME, Harris RC, Bassler BL. 2010. Control of the type 3 secretion system in Vibrio harveyi by quorum sensing through repression of ExsA. Appl Environ Microbiol 76:4996-5004.
4. Paul, P., et al., *Quorum sensing employs a dual regulatory mechanism to repress T3SS gene expression.* mBio, 2025.
5. Chaparian RR, Tran MLN, Miller Conrad LC, Rusch DB, van Kessel JC. 2020. Global H-NS counter-silencing by LuxR activates quorum sensing gene expression. Nucleic Acids Res 48:171-183.
6. Chaparian RR, Olney SG, Hustmyer CM, Rowe-Magnus DA, van Kessel JC. 2016. Integration host factor and LuxR synergistically bind DNA to coactivate quorum-sensing genes in Vibrio harveyi. Mol Microbiol 101:823-40.

7. Cianfanelli, F.R., Cunrath, O. & Bumann, D. Efficient dual-negative selection for bacterial genome editing. *BMC Microbiol* **20**, 129 (2020). https://doi.org/10.1186/s12866-020-01819-2
8. Dalia TN, Chlebek JL, Dalia AB. 2020. A modular chromosomally integrated toolkit for ectopic gene expression in Vibrio cholerae. Sci Rep 10:15398.
